# “TMEM16K is an interorganelle regulator of endosomal sorting”

**DOI:** 10.1101/2020.04.17.046961

**Authors:** Maja Petkovic, Juan Oses-Prieto, Alma Burlingame, Lily Yeh Jan, Yuh Nung Jan

## Abstract

Communication between organelles is essential for their cellular homeostasis. Neurodegeneration reflects the declining ability of neurons to maintain cellular homeostasis over a lifetime^1^. The endolysosomal pathway plays a prominent role in this process by regulating protein and lipid sorting and degradation^2^. Here, we report that TMEM16K, an endoplasmic reticulum lipid scramblase^3^ causative for spinocerebellar ataxia (SCAR10), is an interorganelle regulator of the endolysosomal pathway. We identify endosomal transport as a major functional cluster of TMEM16K in proximity biotinylation proteomics analyses. TMEM16K forms contact sites with endosomes, reconstituting split-GFP with small GTPase RAB7. Our study further implicates TMEM16K lipid scrambling activity in endosomal sorting at these sites. Loss of TMEM16K function led to impaired endosomal retrograde transport and neuromuscular function, one of the symptoms of SCAR10. Thus, TMEM16K-containing ER-endosome contact sites represent clinically relevant platforms for regulating endosomal sorting.

---

Cellular organelles do not act as discrete autonomous units, but rather as interconnected hubs that engage in extensive communication to coordinate their function over a lifetime and maintain cell homeostasis. An emerging theme is that such coordination can be mediated via membrane contact sites (MCS) between distinct organelles. These MCS are specialized microdomains of close proximity between the organelles that modulate essential cellular processes like lipid homeostasis^4–6^ and organelle biogenesis^7^. While multiple human orthologues of MCS proteins^8,9^ have been linked with a broad range of age-related pathologies, the identity and functions of membrane contact sites in maintaining cellular physiology remain an open question.

TMEM16 family of proteins is evolutionarily conserved with members found in all eukaryotes^10^ from amoeba to humans. They are modulators of diverse cellular processes, which when perturbed lead to a variety of genetic disorders^11,12^. Although the TMEM16 family is most frequently associated with their functions as calcium activated chloride channels or phospholipid scramblases, their yeast homolog Ist2 was one of the first reported MCS proteins, playing a vital role in lipid homeostasis at MCS between the ER and plasma membrane^13^. Here we focus on TMEM16K, the mammalian family member most closely related to Ist2 (Supplemental Fig. 1a) and causative of a recessive form of spinocerebellar ataxia (SCAR10)^14^.

We generated mouse models with either ubiquitous or neuron specific loss of TMEM16K (Supplemental Fig. 1b) to evaluate if the pathology is conserved between mouse and human. As impairment of neuromuscular function is a classical symptom of ataxia, we analyzed neuromuscular junctions (NMJ)^15^ in TMEM16K knockout mice at 6 and 24 months of age. Using bungarotoxin staining, we found a progressive reduction in the size of the NMJ (Fig. 1a,b). Moreover, knockout mice displayed increasing hindlimb clasping, a behavioral phenotype marking disease progression in a number of mouse models of neurodegeneration^16,17^ (Fig. 1c, Supplemental Video 1). As TMEM16K is broadly expressed (Supplemental Fig. 1c), we analyzed neuron specific TMEM16K knockout mice and wild type littermates at 24 months of age to evaluate whether loss of TMEM16K in neurons is sufficient to cause the observed phenotypes. These animals lacking neuronal TMEM16K displayed increased hindlimb clasping, as well as an impaired ability to complete a ledge-walking test (Fig. 1d). Together, these results demonstrate a phenotypic linkage between loss of TMEM16K and impaired neuromuscular function that is conserved between mice and human.

**Figure 1.**
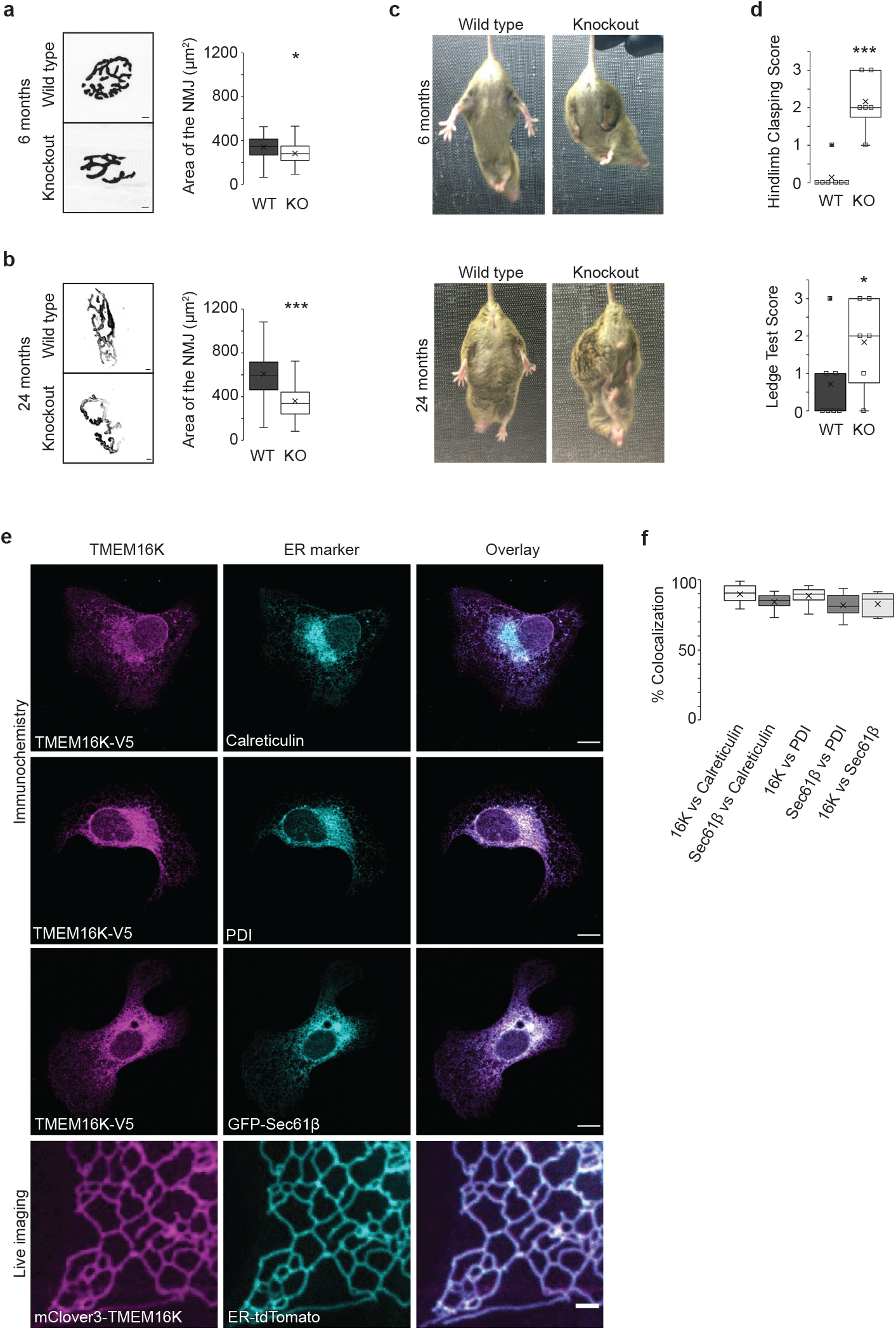
TMEM16K knockout mice display progressive impairment in neuromuscular function. **a.** Representative images and quantification of neuromuscular junction (NMJ) at 6 months of age from wild type (WT, 133 NMJ, 3 animals) and TMEM16K full knockout (KO, 132 NMJ, 4 animals) littermates visualized with the fluorescently labelled α-Bungarotoxin. Scale bar 5 μm, ANOVA, P<0.01* **b.** Representative images and quantification of neuromuscular junction at 24 months of age from WT (137 NMJ, 5 animals) and TMEM16K full KO (126 NMJ, 4 animals) littermates. Scale bar 5 μm, ANOVA, P<0.0001*** **c.** Representative images of hindlimb clasping, of WT and TMEM16K full KO littermates at 6 and 24 months of age. **d.** Quantification of hindlimb clasping and ledge test at 24 months of age of neuron specific TMEM16K KO (6 animals) and their WT (7 animals) littermates P<0.01 *, P<0.0001 *** two-tailed t-test. See Supplemental Video 1. **e. *Rows 1-3:*** Immunocytochemistry of U2OS cells transfected with TMEM16K tagged with V5 epitope and stained for ER-markers Calreticulin, Protein Disulfide-Isomerase (PDI) and Sec61β, respectively. Scale bar 10 μm. **e. *Row 4:*** Snapshot from live imagining in COS7 cells expressing with TMEM16K tagged with mClover3 and ER-tdTomato. Scale bar 2 μm. See Supplemental Video 2. **f.** Quantification of the colocalization of TMEM16K and ER-markers Calreticulin, PDI and Sec61β using Mander’s overlap coefficient, as well as quantification of the colocalization of Sec61β with Calreticulin and PDI measured in the same manner. Sec61β cololicalization with other ER markers is included to provide a meaningful context for the colocalization analysis with TMEM16K, given that Sec61β is a pore forming component of the translocon complex localized exclusively to the ER. (n=38 cells for TMEM16K vs PDI, 44 for TMEM16K vs Calreticulin, 20 for TMEM16K vs Sec61β, 42 for Sec61β vs Calreticulin, 38 for Sec61β vs PDI).

TMEM16K has previously found to be localized to the endoplasmic reticulum^3^, a localization shared with its yeast^18^ and *Drosophila*^19^ homologs. Consistent with these studies, we found TMEM16K localizes to the endoplasmic reticulum (ER) as evident by colocalization with several established endoplasmic reticulum markers (Fig. 1e,f, Supplemental Video 2).

To elucidate the potential cellular functions of TMEM16K in an unbiased manner, we set out to identify the TMEM16K protein interaction network using proximity-dependent biotinylation^20–22^ (BioID). This approach uncovers direct and indirect interactions within a 10-nanometer range of the promiscuous biotinylation enzyme tagged to the protein of interest (Fig. 2a). We tagged TMEM16K with biotin ligase and confirmed retention of both endoplasmic reticulum localization (Supplemental Fig. 2a,b) and biotinylation activity (Supplemental Fig. 2c), permitting the identification of TMEM16K-proximal proteins by mass spectrometry of affinity-purified biotinylated proteins from transfected cells (Fig. 2a). We obtained a list of potential TMEM16K interactors (Fig. 2b, Supplemental Table 1) and, instead of hand-picking a few candidates, we visualized this list as a protein-protein interaction network to identify the most biologically interconnected clusters of proteins which could infer TMEM16 function (Fig. 2b). First, we calculated protein-protein interaction enrichment to determine if the obtained candidate list has more or less interactions among themselves, as compared to a random set of proteins of similar size. Protein-protein interaction enrichment p-value of the TMEM16K network is p<1.0E-16, suggesting biological connection of proteins that interact with TMEM16K. Next, we performed functional enrichment analysis, and overlaid the major functional categories on our candidates, suggesting the presence of functional clusters in our candidate list. Hence, we performed clustering analysis to bioinformatically identify such clusters, defined as highly interconnected nodes, and generated a simplified network of TMEM16K major clusters overlaid with functional enrichment categories (Fig. 2c). As expected, when evaluating a protein over its lifetime, we found clusters involved in protein processing and degradation. Consistent with the function of its *Drosophila* homolog^19^, we also identified a cluster of proteins involved in nuclear organization. Unexpectedly, this analysis revealed that endosomal transport, in particular endosomal retrograde trafficking, is a major cluster in the TMEM16K network (Fig. 2c).

**Figure 2.**
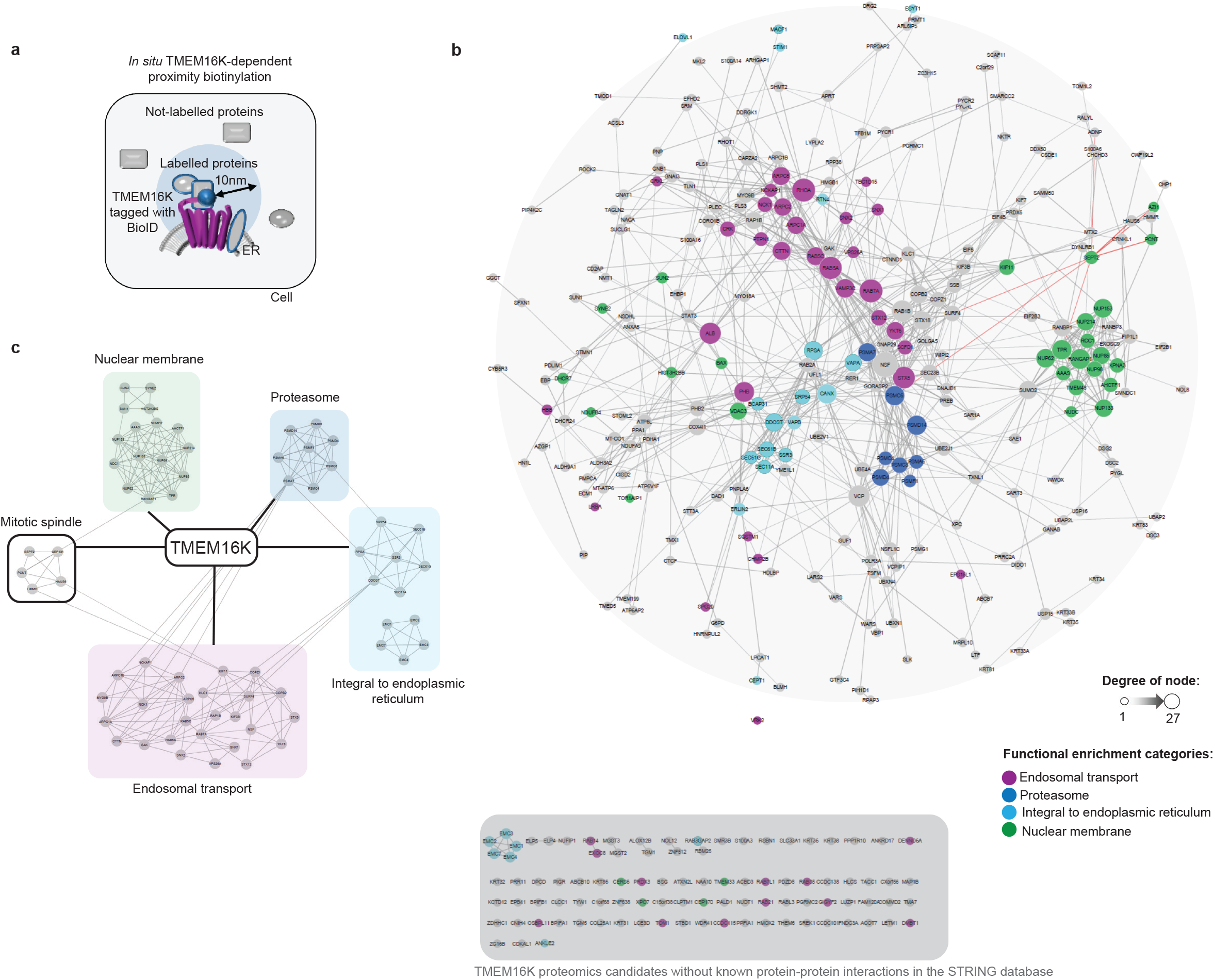
Proteomic mapping of TMEM16K via *in situ* BioID-catalyzed biotin labeling finds endosomal transport as a major functional cluster. **a.** Scheme of proteomic mapping of protein complexes surrounding TMEM16K in the radius of 10 nm via *in situ* proximity biotinylation. **b.** TMEM16K proteome candidate list is visualized as a protein-protein interaction network using the String protein interaction public database in Cytoscape. Candidates without known protein-protein interactions in the String database are represented on the bottom in the gray panel. TMEM16K is omitted from this representation for simplicity. Functional enrichment based on the GO terms was calculated using the String app in the Cytoscape and the major identified categories of functional enrichment were overlayed on our candidates with color-code. *Purple*: Endosomal transport (False Discovery Rate (FDR) p-value 2.49E-4), endosome to Golgi retrograde trafficking (FDR p-value 0.00962); *Cyan:* ER membrane protein complex (FDR 1.5E-5), protein localization to endoplasmic reticulum (FDR p-value 0.00264); *Green*: nuclear membrane (FDR p-value 1.28E-6), nuclear pore (FDR p-value 3.38E-9); *Blue*: proteasome (FDR p-value 5.74E-5). **c.** Bioinformatic analysis of the TMEM16K candidates list with MCODE cluster app in Cytoscape identified major clusters in our dataset, which generated simplified network of TMEM16K proteomics data. Color-coding of functional enrichment analysis was overlaid on the bioinformatically identified clusters. TMEM16K candidate list is provided in Supplemental Table 1.

Dysfunctions of endosomal transport are tightly associated with neurodegenerative diseases^23,24^. As the TMEM16K interactome pointed to endosomal retrograde transport, we investigated whether TMEM16K is required for proper trafficking of the cation-independent mannose-6-phosphate receptor (CI-MPR), which is the best-studied retrograde-transport cargo in mammals^25,26^. In wild type primary mouse embryonic fibroblasts (MEF), when an antibody that recognizes the extracellularly exposed CI-MPR is pulse chased from the plasma membrane, it gets internalized in the endosomes and subsequently transported through the endosomal retrograde pathway to the perinuclear region corresponding to the trans-Golgi network (TGN) within 60 min^1,2^ (Fig. 3a). However, in MEF from TMEM16K knockout mice, the internalized antibody against CI-MPR remained dispersed peripherally during the 60 min period of pulse chase, consistent with a defect in endosome to trans-Golgi retrograde trafficking (Fig. 3a). Reintroduction of TMEM16K rescued this CI-MPR retrograde trafficking defect of mutant MEF (Fig. 3a). As this pathway is also co-opted by a subgroup of pathogens during their entry into cells, we used cholera toxin B (CTxB)^27^ to further corroborate our finding. Indeed, following CTxB internalization, TMEM16K knockout cells displayed reduced CTxB colocalization with the Golgi marker GM130 after 60 min of pulse chase, which can be rescued by reintroduction of TMEM16K (Fig. 3b), confirming that TMEM16K is required for proper endosome to trans-Golgi retrograde trafficking. However, no change was observed in the localization of Golgi complex proteins, nor in the morphology of the Golgi complex (Fig. 3b; Supplemental Fig. 3), suggesting that loss of TMEM16K function does not affect the Golgi complex. Altogether, these data demonstrate that depletion of the ER-resident protein TMEM16K perturbs endosomal retrograde trafficking, a defect similar to that observed upon depletion of known cargo-sorting components^26,28,29^.

**Figure 3.**
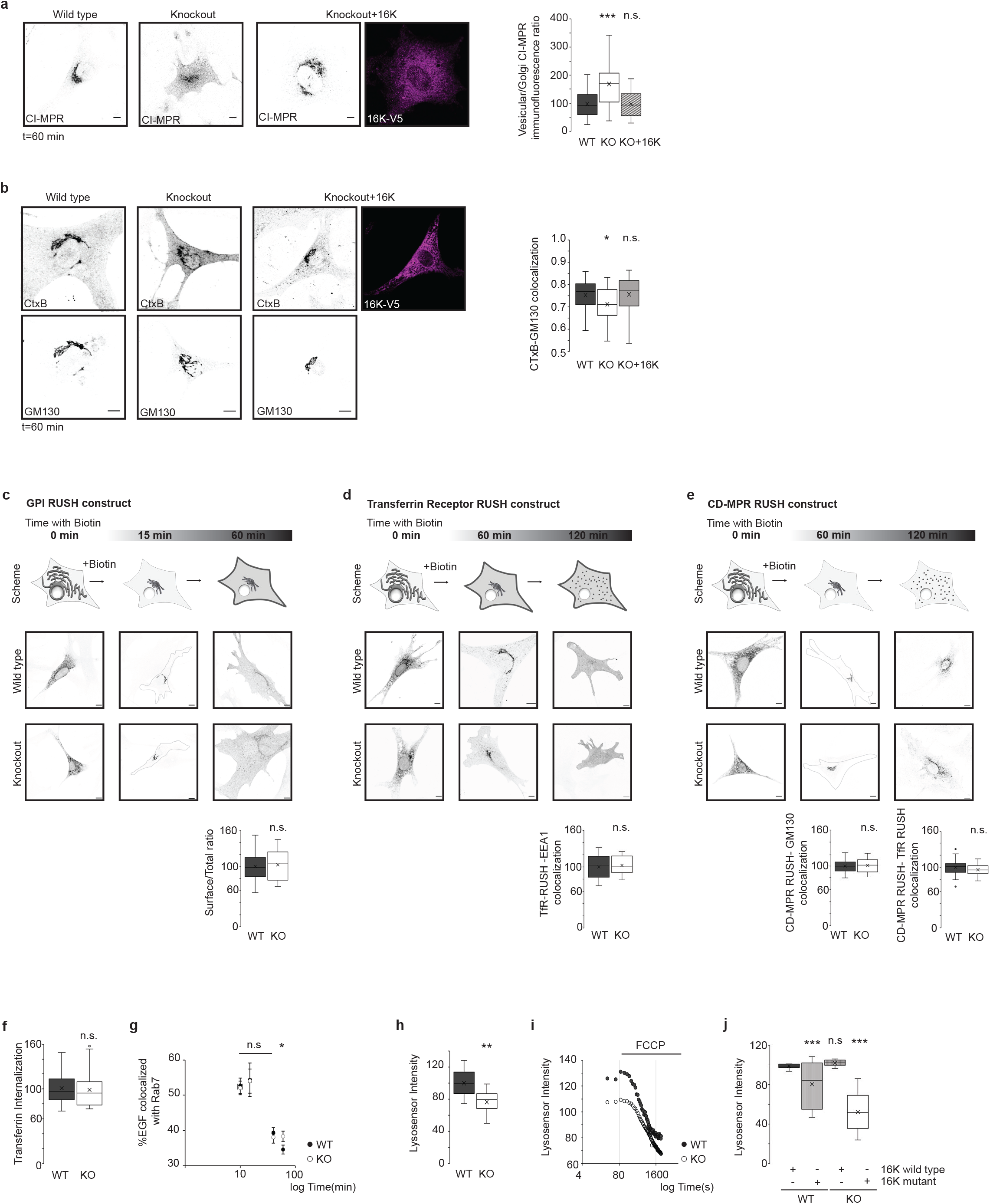
Absence of TMEM16K leads to perturbations of the endosomal sorting. **a. *Left***, Immunofluorescence of pulse chased antibody detecting CI-MPR internalized from the plasma membrane over 60 min time interval in the WT, TMEM16K KO cell and TMEM16K KO cell with reintroduced TMEM16K. Scale bar 10 μm. **a. *Right*,** Ratio of measured intensity between vesicular region of the cell and the region encompassing Golgi (10 × 10 μm^2^). ANOVA, post-test Bonferroni-corrected *t-*test is labelled on the graph with pairwise comparison with WT. P<0.0001 ***. (n=168 for WT, 181 for KO and 134 for KO+16K) **b. *Left*,** Immunofluorescence of Golgi marker GM130 and internalized conjugated cholera toxin B (CtxB) from the plasma membrane over 60 min time interval in the WT, TMEM16K KO cell and TMEM16K KO cell with reintroduced TMEM16K. Scale bar 10 μm. **b. *Right*,** Quantification of the Pearson’s correlation coefficient measuring colocalization of GM130 and CtxB. ANOVA with post-test Bonferroni-corrected *t-*test is labelled on the graph with pairwise comparison with WT, P<0.01 *. (n= 40 for WT, 57 for KO, and 56 for KO+16K) **c-e.** RUSH assay. WT and TMEM16K KO cells are transfected with RUSH constructs, where marker proteins are blocked in the endoplasmic reticulum and released upon addition of biotin enabling monitoring of their transport through biosynthetic pathway. **c.** RUSH assay. Scheme of mCherry-GPI RUSH construct biosynthetic pathway, with representative images of WT and TMEM16K KO cells expressing mCherry-GPI RUSH construct at 0, 15 and 60 min after addition of biotin, and quantification of surface vs. total mCherry-GPI RUSH construct at 60 min after biotin addition. P>0.01 n.s. two-tailed Student t-test. (n= 53 for WT, 56 for TMEM16K KO) **d.** RUSH assay. Scheme of mCherry-TfR RUSH construct biosynthetic pathway, with representative images of WT and TMEM16K KO cells expressing mCherry-TfR RUSH construct at 0, 60 and 120 min after addition of biotin and quantification of the Pearson’s correlation coefficient measuring colocalization of mCherry-Tfr RUSH construct at 120 min after biotin addition with EEA1. P>0.01 n.s. two-tailed Student t-test. (n=129 for WT, 111 for TMEM16K KO) **e.** RUSH assay. Scheme of GFP-CD-MPR RUSH construct biosynthetic pathway, with representative images of WT and TMEM16K KO cells expressing GFP-CD-MPR RUSH construct at 0, 60 and 120 min after addition of biotin. This experiment is quantified at time points 60 and 120 min after addition of biotin: quantification of the Pearson’s correlation coefficient measuring colocalization of GFP-CD-MPR RUSH construct at 60 min after biotin addition with GM130. (n=126 for WT, 127 for TMEM16K KO), and quantification of the Pearson’s correlation coefficient measuring colocalization of GFP-CD-MPR and mCherry-TfR RUSH constructs at 120 min after biotin addition. (n=144 for WT, 134 for TMEM16K KO), P>0.01 n.s. two-tailed Student t-test. **f.** Quantification of fluorescence intensity of internalized fluorescently conjugated transferrin (60 min) in the WT (100) and TMEM16K KO (50) cells, P>0.01 n.s. two-tailed t-test. **g.** EGF-Alexa647 pulse-chase experiment was quantified for colocalization with endogenous Rab7, P<0.05 * two-tailed t-test between WT and KO at each measured time point (n=130 for WT and 117 for KO at 10min, 91 for WT and 103 for KO at 15min, 77 for WT and 61 for KO at 40 min, 89 for WT and 118 for KO at 60 min). **h.** Quantification of fluorescence intensity of Lysosensor Green DNP-189 in WT and TMEM16K KO as a measure of acidification. ANOVA P<0.001 ** (n=114 for WT and 116 for KO). **i.** Representative trace from 3 independent experiments of protonophore FCCP at a final concentration 2 μM added at 120 seconds to cells loaded with Lysosensor Green DNP-189. The slope following FCCP was less steep indicative of slower rate of proton leak as quantified by linear equation (WT slope is −0.0445, y=-0.0445x+128.45, R^2^=0.9537; KO slope is −0.0305, y=−0.0305x+109.5, R^2^=0.9581) **j.** Evaluation of wild type and mutant TMEM16K cDNA ability to rescue acidification defect. WT or TMEM16K KO cells were co-transfected with mCherry-CAAX to visualize transfected cells, and TMEM16K wild type cDNA (TMEM16K-FLAG) or TMEM16K mutant cDNA (Ca5MUT-FLAG) and evaluated for acidification with Lysosensor Green D-189. ANOVA with post-test Bonferroni-corrected *t-*test is labelled on the graph with pairwise comparison with WT+ 16K wild type, P<0.0001 *** (3 biological replicates, n=WT+ 16K wild type 50; WT + 16K mutant 50; KO+ 16K wild type 40; KO+ 16K mutant 40 cells).

To ensure that the observed defect in endosomal retrograde transport is not due to impaired anterograde secretory pathway, we have taken advantage of a novel approach that allows synchronization of protein transport through the biosynthetic pathway^30,31^. Using this RUSH system we tracked the biosynthetic transport of three transmembrane proteins with different steady state distributions: the glycosylphosphatidylinositol anchor (GPI; transported to plasma membrane), the transferrin receptor (TfR; transported to plasma membrane, early endosomes, and recycling endosomes), and the cation-dependent mannose-6-phosphate receptor (CD-MPR; transported from TGN directly to early/late endosomes). We found no difference in the transport through the biosynthetic pathway between TMEM16K wild type and knockout cells (Fig. 3c-e), showing that the anterograde secretory pathway is unaffected.

To look for evidence for potential defects upstream of the endosomal retrograde sorting, we evaluated first whether endocytosis is affected by examining the internalization of fluorescently labelled transferrin. We found no differences between TMEM16K wild type and knockout cells (Fig. 3f). Next, we performed EGF pulse-chase experiments to further evaluate maturation from early to late endosomes. We found no difference between the wild type and knockout cells in the colocalization of EGF with the late endosomal marker Rab7 at the 10 min, 15 min and 40 min time points (Fig. 3g), indicating that the mutant phenotype arose from a defect at or after the Rab7 stage of endolysosomal maturation. These results show that the upstream endosomal pathway is unaffected in TMEM16K knockouts. However, at 60 min there was a larger fraction of EGF that was retained in Rab7 endosomes in TMEM16K knockout cells, compared to wild type cells (Fig. 3g), suggesting defect in endosomal sorting.

To evaluate whether endolysosomal pathway downstream of endosomal sorting was perturbed, we examined a major distinguishing feature of endolysosomal maturation, acidification. Using Lysosensor Green DNP-189, a fluorescent pH indicator that partitions into acidic organelles, we found that acidification is impaired in the absence of TMEM16K (Fig. 3h). To confirm that the differences in DNP-189 fluorescence reflected differences in pH within organelles, we utilized the protonophore FCCP to selectively eliminate the pH gradient. Consistent with an inability to form and/or maintain a proton gradient, TMEM16K knockout cells displayed both a slower rate of FCCP-induced proton leak and an impaired ability to stabilize proton loss compared with wild type cells (Fig. 3i). To further test the TMEM16K involvement in the observed defect, we performed rescue experiments with wild type TMEM16K or mutant TMEM16K with substitutions of the conserved calcium binding acidic residues required for protein function (E448Q/D497N/E500Q/E529Q/D533N)^3,32,33^. We expressed wild type or mutant TMEM16K in primary cells from wild type or TMEM16K knockout mice. Transfecting mutant TMEM16K into wild type cells has dominant negative effect. Expression of wild type, but not mutant TMEM16K in primary cells lacking TMEM16K rescued the acidification defect, demonstrating that TMEM16K is required for normal maturation of endolysosomal compartments (Fig. 3j). Taken together, these results show that loss of TMEM16K causes a defect in endosomal retrograde sorting, and deficiencies within the later stages of the endolysosomal system.

The requirement of TMEM16K for endosomal retrograde trafficking raised the question how this endoplasmic reticulum localized protein affects endosomes. Given that proteomics revealed that TMEM16K is in the proximity of the endosomal compartment for direct or indirect interactions, we considered the possibility that TMEM16K facilitates endosomal sorting through interorganelle communication at the sites of contact between the ER and endosomes. Membrane contact sites between the ER and endosomes have been shown to increase as endosomes mature^34^, to define endosome fission^7^, and to control association of endosomes with the cytoskeleton^35^, all of which are essential for proper endolysosomal function. The TMEM16K proteomics dataset contained multiple proteins known to function at ER-endosomal contact sites including VAPA and VAPB^35–37^, SNX1 and SNX2^35^, Rab7A^34,37,38^, PTP1B^39^ (Fig. 2b; Supplemental Table 1), suggesting that TMEM16K acts at or in the proximity of these membrane contact sites.

Hence, we applied in cell culture the same proximity biotinylation approach labeling within a 10-nanometer range used for proteomics, in order to visualize TMEM16K-dependent labeling of Rab7-positive endosomes (Fig. 4a). We found that TMEM16K-proximity dependent labeling overlapped with endogenous Rab7. For a dynamic view, we performed live imaging of fluorescently labeled TMEM16K in the ER, with fluorescently labeled Rab7, and fluorescently conjugated EGF in endosomes (Supplemental Video 3, Fig. 4b). These experiments revealed highly dynamic movements of both compartments, as well as their contacts. Using structured illumination microscopy (SIM), we imaged TMEM16K and Rab7-positive endosomes, and visualized dually labeled ER-endosome contact sites (Fig. 4c-e). Next, we used the split-GFP system to specifically evaluate TMEM16K interorganelle contact sites. Split-GFP reconstitution has been extensively used to detect interorganelle contact sites^37–39^. TMEM16K was tagged with a GFP_11_ fragment, while several ER (VAPA, OSBPL8) and endosomal (Rab7, OSBPL9, OSBPL11, VPS26, VPS35, SNX1, SNX2) proteins were tagged with the GFP_1-10_ fragment. We selected proteins that were identified in the TMEM16K proteomics and implicated in ER-endosomal MCS (VAPA, Rab7, OSBPL11, VPS26, SNX1, SNX2), as well as proteins that are not TMEM16K proteomics interaction candidates, but are known to participate in similar processes/compartments (OSBPL8, OSBPL9, VPS35) as negative controls. Since TMEM16K forms a dimer^3,32^, we validated the split-GFP approach by expressing TMEM16K-GFP_1-10_ and GFP_11_-TMEM16K to reconstitute the split-GFP (Fig. 4f). Reconstitution of split-GFP between TMEM16K and any of the tested candidates suggests that they directly interact, bringing the two GFP fragments to such close proximity that they can reconstitute the fluorescent GFP. Inability to reconstitute split-GFP suggests that TMEM16K and the tested candidate are not in close proximity, though we cannot exclude the possibility that steric hindrance may prevent the split-GFP reconstitution (Fig. 4g). Out of all the combinations tested, only Rab7 reconstituted split-GFP with TMEM16K, demonstrating that ER-localized TMEM16K forms contacts with Rab7 endosomes (Fig. 4h; Supplemental Fig. 4). Rab7 is a GTPase that cycles between inactive GDP bound states and active GTP bound states. To evaluate further the specificity of TMEM16K interaction with the Rab7, we generated Rab7 mutants: constitutively active Q67L that mimics permanently GTP-bound Rab7 and inactive T22N that mimics permanently GDP-bound Rab7^43^. We found that TMEM16K was able to reconstitute split-GFP only with the constitutively active Rab7 Q67L mutant (Fig. 4i), but not with the inactive Rab7 T22N mutant (Fig. 4j), further validating the specificity of the observed contact between TMEM16K and Rab7 endosomes.

**Figure 4.**
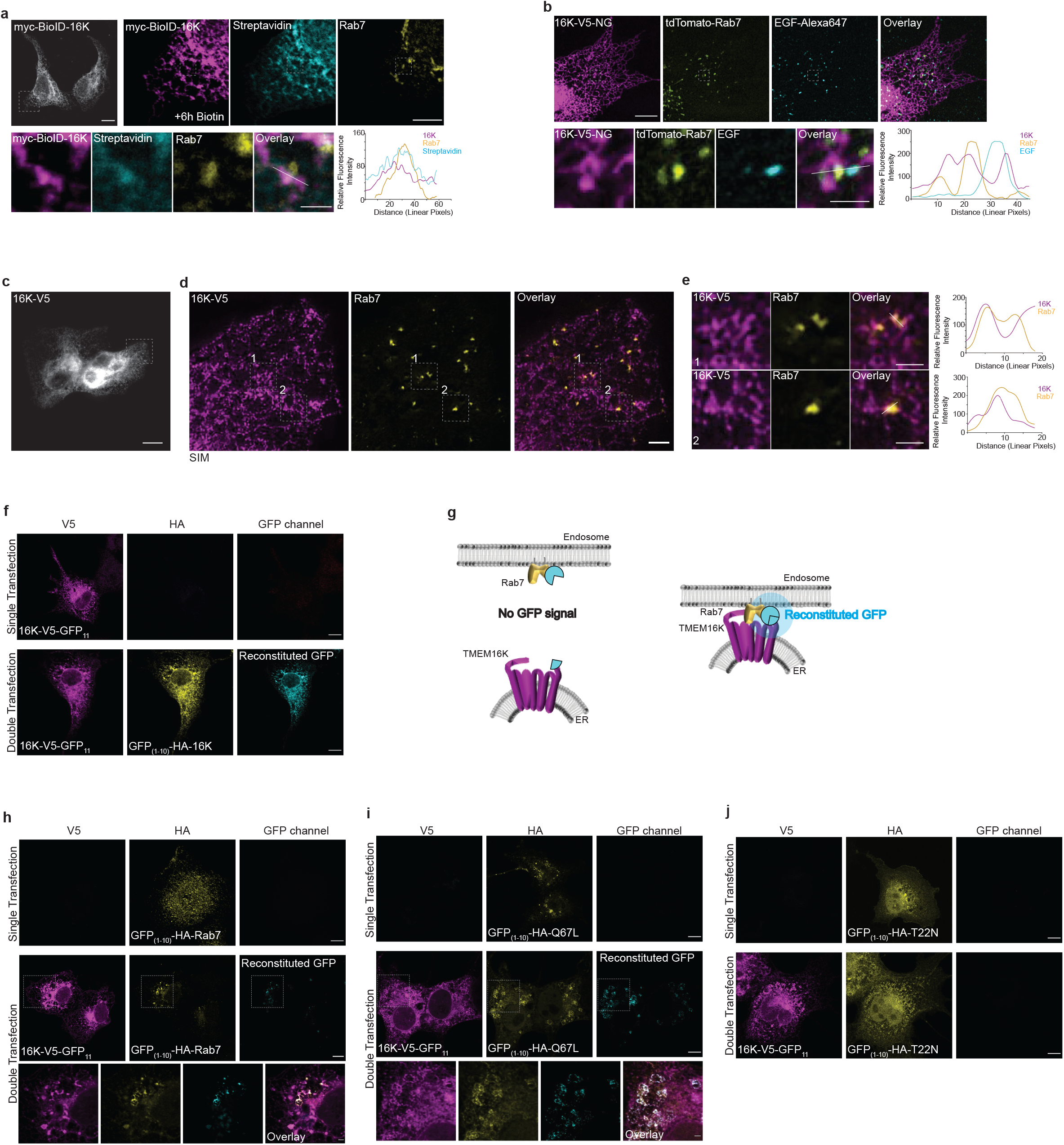
TMEM16K at ER-endosome membrane contact sites. **a.** Immunocytochemistry to visualize TMEM16K proximity labeling of endosomes. COS7 cells were transfected with TMEM16K tagged with proximity biotinylation enzyme, incubated with biotin for 6 h and immunostained with fluorescently conjugated Streptavidin and antibody against endogenous Rab7. **a. *Row 1, Left:*** View of the entire cell expressing myc-BioID-TMEM16K. Scale bar 10 μm. **a. *Row 1, Right:*** Magnified region of the cell showing myc-BioID-TMEM16K, its pattern of proximity biotinylation and endogenous Rab7. Scale bar 5 μm. **a. *Row 2:*** High magnification insets with line scan quantification of the three channels marked in overlay. Scale bar 1 μm. **b.** Live imaging of the U2OS cells transfected with TMEM16K-V5-mNeonGreen (TMEM16K-V5-mNG), tdTomato-Rab7, and EGF-Alexa647, pulse chased for 45 min, imaged with spinning disk confocal microscope. See Supplemental Video 3. **b. *Row 1:*** Snapshots of the live imaging showing TMEM16K, Rab7, EGF and their overlay. Scale bar 10 μm. **b. *Row 2***, High magnification insets with line scan quantification of the three channels marked in overlay. Scale bar 0.5 μm **c.** Widefield image to view entire cell expressing TMEM16K-V5. Scale bar 10 μm. Inset marks cell region imaged with structured illumination microscopy (SIM). **d.** Single plane structured illumination microscopy of U2OS cells transfected with TMEM16K-V5 and immunolabelled for endogenous Rab7. Scale bar 2 μm. **e.** High magnification insets 1 and 2 from SIM images with corresponding line scan quantification of the two channels marked in overlay. Scale bar 1 μm. **f.-j.** Split-GFP assay. Immunocytochemistry of single and double transfected cells with TMEM16K-V5-GFP_11_ and GFP_(1-10)_ tagged TMEM16K, wild type Rab7 or Rab7 mutants. Single transfected cells were used as control, as reconstituted split-GFP signal can only be present in double transfected cells. **f.** Split-GFP assay positive control with cells transfected with TMEM16K-V5-GFP and GFP_(1-10)_-HA-TMEM16K, as TMEM16K acts as dimer. Scale bar 10 μm. **g.** Scheme of the split-GFP experiment with Rab7 where molecule of the GFP can be reconstituted only if the two proteins contact **h.** TMEM16K-V5-GFP and GFP_(1-10)_-HA-Rab7 is the only tested combination able to reconstitute the split-GFP. Scale bar 10 μm, insets 2 μm. See Supplemental Figure 3. for all the tested proteins and negative controls. **i.** Split-GFP reconstitution assay evaluating TMEM16K-V5-GFP_11_ ability to reconstitute GFP with constitutively active mutant of Rab7 Q67L tagged with GFP_(1-10)_ Scale bar 10 μm, inset 2 μm. **j.** Split-GFP reconstitution assay evaluating TMEM16K-V5-GFP_11_ ability to reconstitute GFP with inactive mutant of Rab7 T22N tagged with GFP_(1-10)_ Scale bar 10 μm.

The yeast homolog Ist2 mediates ER-plasma membrane contact sites by directly binding plasma membrane specific phosphatidylinositol-(4, 5)-bisphosphate (PtdIns(4,5)P_2_) with its C-terminus^44^. The presence of a series of positively charged residues in the TMEM16K N-terminal cytosolic domain prompted us to hypothesize that, in addition to its interaction with endosomal proteins like Rab7, TMEM16K may directly bind phosphatidylinositols specific for the endolysosomal compartments. To test this hypothesis, we purified the N-terminal domains of TMEM16K and two other mammalian family members as control, TMEM16F and TMEM16A, to evaluate their lipid binding via a protein lipid overlay assay (Fig. 5a,b,c). We found that the N-terminal domain of TMEM16F binds plasphosphatidylinositol-(3,4,5)-phosphate (PtdIns(3,4,5)P_3_), as recently reported^45^. Unlike TMEM16F and TMEM16A, TMEM16K specifically bound phosphatidylinositols present in endolysosomal compartments, including phosphatidylinositol-3-phosphate (PtdIns3P) (Fig. 5c), the major phosphatidylinositol in endosomes. To evaluate the functional requirement of the N-terminal domain, we generated N-terminal deletion mutant of TMEM16K (Δ1-169 amino acids) (Fig. 5d). Whereas we used the N-terminal 255 amino acids in the protein overlay assay, mutation of the 171th amino acid is causative for human pathology, so we tested for a truncation mutant that retains this residue. This TMEM16K truncation mutant with N-terminal deletion properly localized to endoplasmic reticulum and could still reconstitute split-GFP with Rab7, demonstrating that the N-terminal domain is not required for TMEM16K contacts with endosomes (Fig. 5e). However, with N-terminal deletion the TMEM16K truncation mutant was not able to rescue endosomal retrograde transport defect of cells from TMEM16K knockout mice, showing that the N-terminal domain is required for TMEM16K function (Fig 5f,g). This functional requirement is reminiscent of the functional requirement of the binding of the TMEM16F N-terminal domain to plasma membrane phosphatidylinsoitols for the regulation of TMEM16F gating^45^. Our findings suggest that the ER-localized TMEM16K forms contact sites with endosomes, where it binds active GTP-bound Rab7 and endolysosomal phosphatidylinositols like PtdIns3P (Fig. 5f).

**Figure 5.**
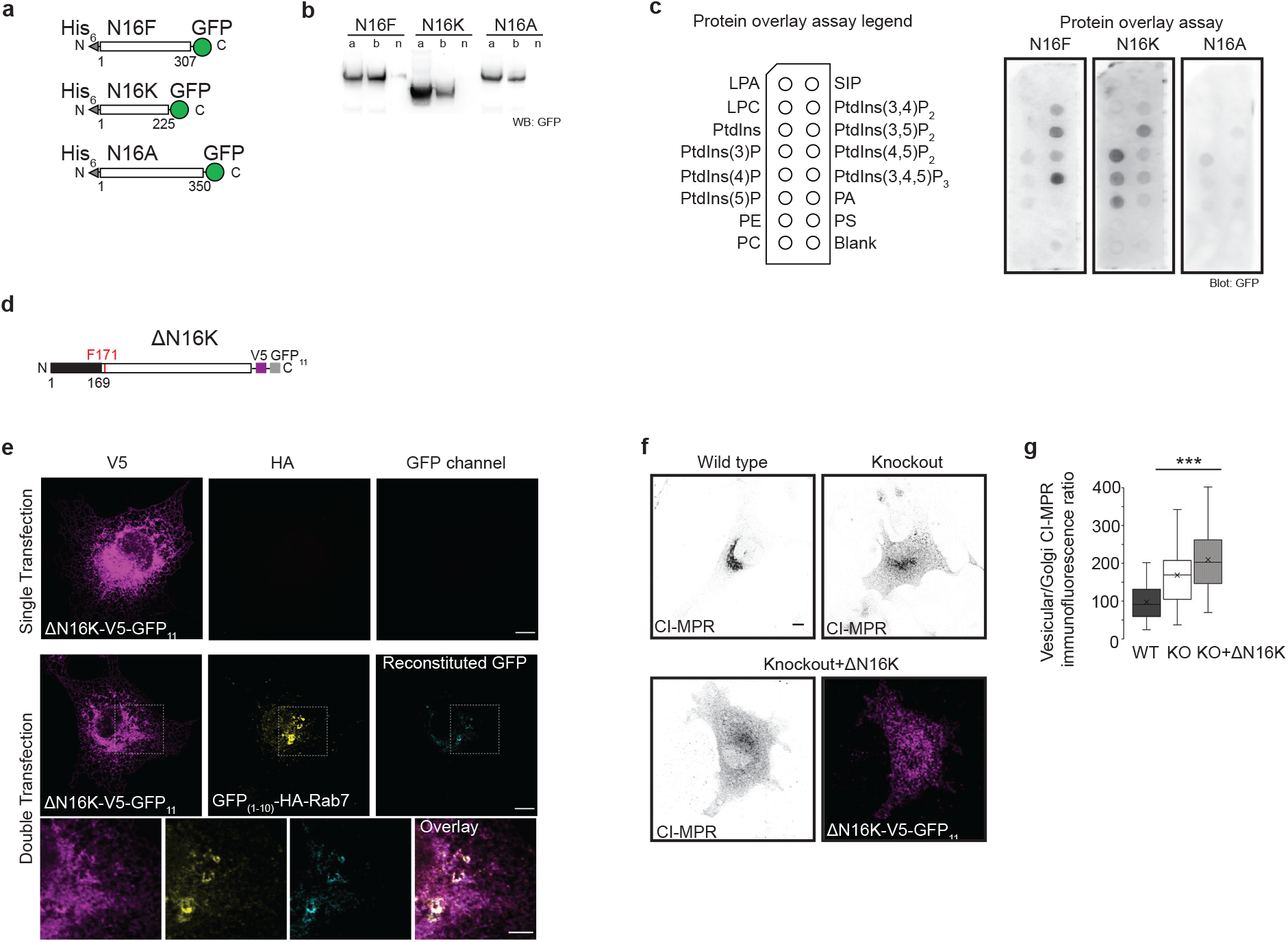
TMEM16K and phosphatidylinositols. **a.** Scheme of purification constructs of N-terminal cytosolic domains of TMEM16F (N16F), TMEM16K (N16K) and TMEM16A (N16A) **b.** Western blot of purified N-terminal cytosolic domains (a- after dialysis, b- before dialysis, n- unbound) revealed with anti-GFP antibody and horseradish peroxidase. **c.** Lipid binding of predicted phosphatidylinositol binding domains was evaluated by loading 10 μg of purified proteins to the PIP-strips and visualized with blotting with anti-GFP antibody and horseradish peroxidase. Representative images of 3 independent blots. **d.** Scheme of N-terminal truncation mutant of TMEM16K where 1-169 amino acids were deleted. **e.** Split-GFP reconstitution assay to evaluate N-terminal truncation mutant for its ability to reconstitute GFP with Rab7. Cell were single or double transfected cells with ΔN-terminal TMEM16K-V5-GFP_11_ and GFP_(1-10)_ tagged Rab7, Scale bar 10 μm for rows 1, 2, and 5 μm for inset in the row 3. **f.** Representative images of the ability of ΔN16K-V5-GFP_11_ to rescue endosomal retrograde trafficking defect when introduced in TMEM16K KO cells as measured by CI-MPR assay described in Fig. 3a(WT and KO from Fig. 3a repeated for clarity of comparison) Scale bar 10 μm. **g.** Quantification (same data from Fig. 3a added to this graph for clarity of comparison). ANOVA with post-test Bonferroni-corrected *t-*test is labelled on the graph with pairwise comparison with WT, P<0.0001 *** (n= 161 cells KO + ΔN16K-V5-GFP_11_).

Next, we sought to address how TMEM16K regulates endosomal function. Like TMEM16K, the VAPA and VAPB proteins, which were detected in the TMEM16K proteomics, act at ER-endosome membrane contact sites and can affect endosomal retrograde trafficking; VAPA and VAPB regulate PtdInsI4P levels on endosomes and subsequent WASH-dependent actin nucleation^35^. If TMEM16K indirectly affects endosomal sorting through this VAPA/B pathway, we would expect it to be associated with similar cellular defects. We first evaluated whether ER-endosome membrane contact sites are globally perturbed in the absence of the TMEM16K. Using electron microscopy^46^, we found no difference in the percentage of endosomes in close proximity (>30 nm) to ER between wild type and TMEM16K knockout cells (Supplemental Fig. 5a,b), consistent with the presence of multiple proteins maintaining these contacts^47,48^. We have further strengthened these observations with proximity ligation assay (PLA) *in situ*, a powerful novel approach to study contact sites alterations in a quantitative manner. Using VAPB and Rab7 as markers of ER and endosomes, respectively, we found no difference in the extent of ER-endosome MCS as measured by the number of PLA puncta between the wild type and TMEM16K knockout cells (Supplemental Fig. 5c,d), corroborating that the absence of TMEM16K does not globally perturb ER-endosome MCS. As cells lacking VAPA/B accumulate PtdIns4P on endosomes^35^ we next assessed PtdIns4P distribution by using the reporter GFP-P4M SidM^49^, and found no perturbation in the absence of TMEM16K (Figure 6a). Likewise, there was no alteration of plasma membrane PtdIns(4,5)P_2_ visualized with the 2PH-PLCΔ-GFP biosensor or anti-PIP2 antibody (Supplemental Fig. 3). Furthermore, unlike VAPA/B DKO cells^35^, we found no perturbation of the actin cytoskeleton in cells lacking TMEM16K (Fig. 6a). However, by utilizing P40PX-EGFP to detect the distribution of PtdIns3P^50^, we found enlarged PtdIns3P vesicles in the absence of TMEM16K (Fig. 6b,d). Reintroducing TMEM16K in KO cells fully rescued the enlarged PtdIns3P vesicles phenotype (Fig. 6b,d). PtdIns3P is a precursor for the generation of PtdIns(3,5)P_2_ by the only mammalian PtdIns 5-kinase, PIKfyve^51^, which when perturbed also leads to enlarged endosomes. Thus, we wondered if this conversion is perturbed in the absence of TMEM16K. As we were not able to reliably visualize PtdIns(3,5)P_2_ with fluorescent reporter in primary cells from wild type and TMEM16K knockout mice, we used the pharmacological inhibitor of PIKfyve^52^, YM201636, and visualized its effects on PtdIns3P (Fig. 6c). Inhibiting PIKfyve in TMEM16K wild type cells recapitulated the TMEM16K knockout phenotype. However, in the TMEM16K knockout cells we did not observe additional cumulative effect, suggesting that conversion of PtdIns3P to PtdIns(3,5)P_2_ is impaired in the absence of TMEM16K (Fig. 6c,d). Altogether, our results strongly indicate that, while there could be coordination of TMEM16K and VAPA/B MCS functions in mediating endosomal retrograde trafficking, TMEM16K affects endosomal sorting in a manner independent of the VAPA/B pathway.

**Figure 6.**
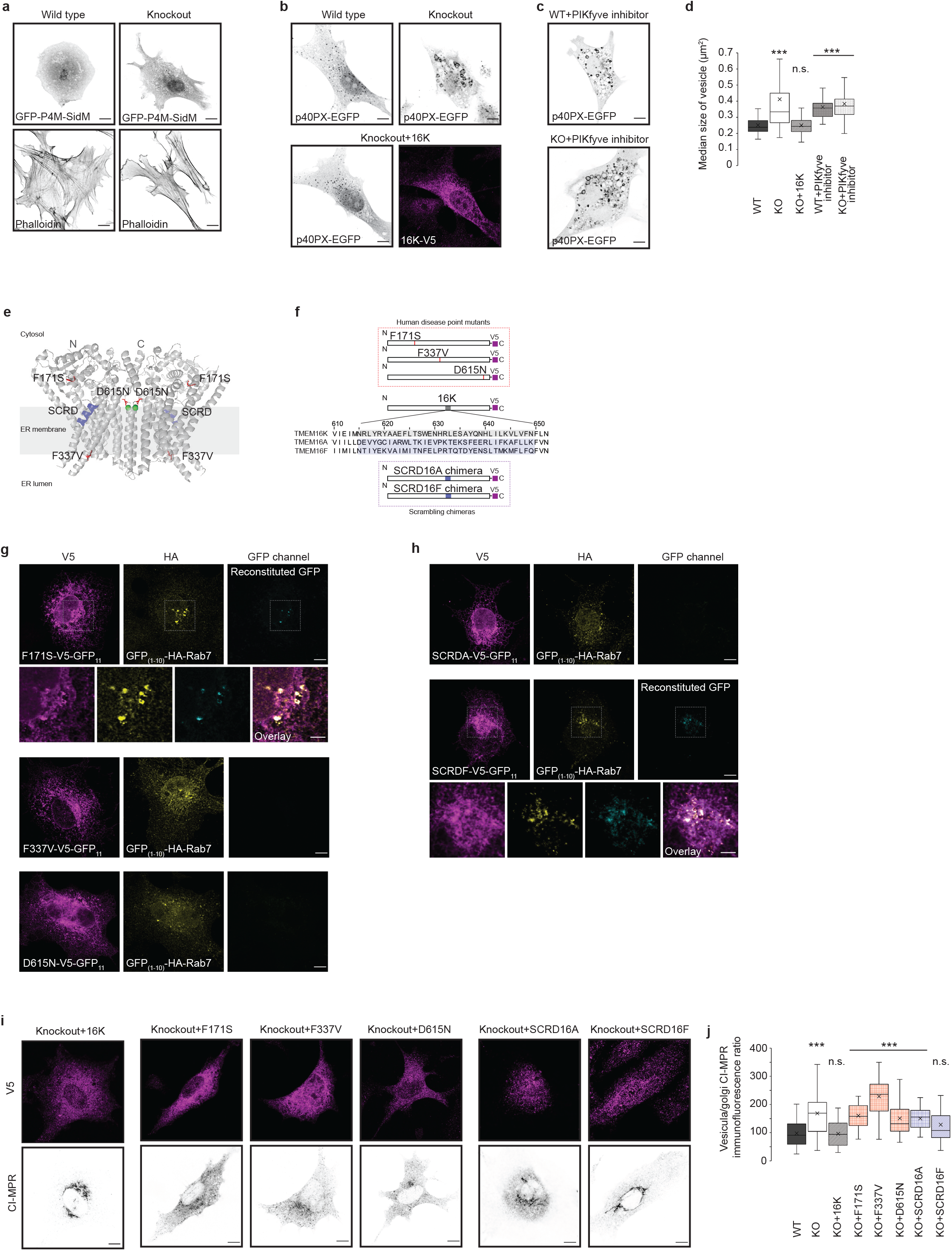
TMEM16K activity is required for endosomal retrograde trafficking. **a.** Confocal images of WT and TMEM16K KO cells transfected with GFP-P4M-SidM, biosensor for PtdIns4P or labelled with fluorescence conjugated phalloidin which specifically labels actin network. Representative images from 2 independent experiments. Scale bar 10 μm. **b.** Confocal images of WT, TMEM16K KO cells and KO cells with reintroduced TMEM16K cells transfected with PtdIns3P biosensor P40PX-EGFP. Scale bar 10 μm. **c.** Confocal images of WT and TMEM16K KO cells transfected with PtdIns3P biosensor P40PX-EGFP and treated with PIKfyve kinase inhibitor YM201636. Scale bar 10 μm. **d.** Quantification of the median size of the PtdIns3P positive vesicles per cell visualized with P40PX-EGFP in the WT (n=131 cells), TMEM16K KO (131), and TMEM16K KO cells transfected with wild type TMEM16K cDNA (153), as well as WT (150) and TMEM16K KO (156) cell treated with PIKfyve inhibitor YM201636. ANOVA with post-test Bonferroni-corrected *t-*test is labelled on the graph with pairwise comparison with WT. P>0.01 n.s. P<0.0001 ***. **e.** Topology of TMEM16K dimer in the endoplasmic reticulum membrane with labeled human disease point mutations and SCRD (minimal lipid scrambling domain) based on published crystal structure^3^ and represented in Pymol. **f.** Scheme of the human disease point mutation constructs and SCRD chimeras where TMEM16K SCRD homology domain was replaced with corresponding 35 aa from non-scramblase TMEM16A or scramblase TMEM16F. **g.** Split-GFP reconstitution assay. Human point mutants (F171S, F337V, D615N) tagged with GFP_11_ were evaluated for their ability to reconstitute GFP when co-transfected with GFP_(1-10)_ tagged Rab7. Scale bar 10 μm, with higher magnification inset 5 μm. **h.** Split-GFP reconstitution assay. Scrambling domain chimeras (SCRDA, SCRDF) tagged with GFP_11_ were evaluated for their ability to reconstitute GFP when co-transfected with GFP_(1-10)_ tagged Rab7. Scale bar 10 μm, with higher magnification inset 5 μm. **i.** Representative images of the ability of human disease point mutants and SCRD chimeras to rescue endosomal retrograde trafficking defect when introduced in TMEM16K KO (KO+ 16K cDNA from Fig. 3a repeated for clarity of comparison) Scale bar 10 μm. **j.** Quantification as done in Fig. 3a (same data added to this graph for clarity of comparison). ANOVA with post-test Bonferroni-corrected *t-*test is labelled on the graph with pairwise comparison with WT, P>0.01 n.s., P<0.0001 *** (n= KO+F171S 145, KO+F337V 138, KO+ D615N 155, KO+ 16KSCRD16A 130, KO+ 16KSCRD16F 145 cells).

To better define human mutations linked to spinocerebellar ataxia (SCAR10), we tested three known single amino acids missense mutations (Phe171Ser, Phe337Val and Asp615Asn; Fig. 6e,f)^3^. These human disease variants expressed normally and localized to the endoplasmic reticulum in heterologous cells (Supplemental Fig. 6). Out of these three, Phe171Ser was able to reconstitute split-GFP with Rab7 (Fig. 6g). Remarkably, unlike wild type TMEM16K that was able to completely rescue the CI-MPR retrograde trafficking defect in TMEM16K knockout cells, these three human disease variants were unable to rescue (Fig. 6i,j). Thus, our results suggest that impairment of the endosomal pathway could be a contributing factor in the development of the SCAR10 pathology.

Given that some TMEM16K mutants can reconstitute split-GFP with Rab7, but are unable to rescue the CI-MPR retrograde trafficking defect in TMEM16K knockout cells (Fig. 5d,e, Fig. 6g,I,j), we conclude that proximity to endosomes is required, but not sufficient for TMEM16K cellular function. TMEM16K was recently demonstrated to possess calcium regulated phospholipid scramblase activity^3,53,54^, translocating phospholipids bidirectionally down their concentration gradients. Therefore, we wondered whether TMEM16K-scrambling function is required for endosomal sorting. Grafting the 35 amino acids constituting the minimal scrambling domain (SCRD)^55^ of the TMEM16F scramblase (Fig. 6e,f) onto the TMEM16A calcium-activated chloride channel conveyed scrambling activity to the chimera. Similarly, grafting SCRD homology region of TMEM16K onto TMEM16A conveyed scrambling activity to the chimera, while grafting SCRD homology region of non-scramblase family members did not convert TMEM16A into a scramblase^53^. To evaluate whether TMEM16K scrambling function is required for endosomal retrograde trafficking, we used the same established approach and generated chimeras substituting the minimal scrambling domain (SCRD) of TMEM16K with that of non-scramblase TMEM16A or scramblase TMEM16F (Fig. 6e,f). Both TMEM16K-SCRD16A and TMEM16K-SCRD16F chimeras can be efficiently expressed and correctly localized to endoplasmic reticulum (Supplemental Fig. 6). When reintroduced into TMEM16K KO cells, the putative non-scramblase chimera TMEM16K-SCRD16A failed to reconstitute split-GFP with Rab7 (Fig 6h) and it could not rescue the retrograde trafficking defect as assayed by CI-MPR internalized antibody distribution (Fig. 6i,j). However the putative scramblase chimera TMEM16K-SCRD16F was able to reconstitute split-GFP with Rab7 (Fig. 6h) and rescue the endosomal retrograde trafficking defect (Fig. 6i,j), suggesting that the lipid scrambling activity of TMEM16K could be required for endosomal sorting.

In conclusion, our study has discovered that TMEM16K acts as an interorganelle regulator of ER-endosomal communication at their sites of contact. Loss of TMEM16K leads to dysfunction of the endolysosomal pathway, which could be rescued by wild type TMEM16K but not mutant TMEM16K bearing the human disease point mutations causing spinocerebellar ataxia. In addition to the observed cellular defects, we found progressive deterioration of the neuromuscular function in the TMEM16K knockout mice. Indeed, dysfunctions in endosomal sorting are known to accumulate over time and represent a converging mechanism shared by a broad range of neurodegenerative diseasess^23,56^. The defect in the later stages of the endolysosomal pathway we observed could be caused by mistargeting of proteins needed for the late endosome/lysosome function due to defects in endosome sorting in the absence of TMEM16K.

Our results are consistent with a model in which the TMEM16K phospholipid scrambling function could, upon TMEM16K activation by binding of the specific phosphatidylinositols and calcium^45^, selectively modulate the local lipid environment in the endoplasmic reticulum at these sites of contact. Distribution of phospholipids across the ER membrane is thought to be largely symmetrical, with the exception of phosphatidylserine^54,57^, and TMEM16K was shown to be required for its calcium-induced leaflet redistribution^54^. A number of recent studies have highlighted the importance of lipid microdomains in protein sorting^58^. For example, translocation of phosphatidylserine across leaflets is required for sorting at the trans-Golgi complex in yeast^59^, and loss of phosphatidylserine asymmetry impairs sorting of early endosomes in *C. elegans*^60^. Just how modulating leaflet composition in the ER would affect endosomal sorting is an interesting open question. TMEM16K-modulated lipid availability in the ER could locally recruit proteins^61^, modulate local protein activity, or promote direct transfer of lipids between ER and endosomes at the sites of contact, providing necessary cues for endosomal sorting. Interestingly, the TMEM16K proteomics includes OSBP-related protein 11 (OSBPL11), lipid transfer proteins reported to localize at the Golgi-late endosome interface^62^. Altogether, our results open the possibility that endosomal sorting could be enabled by modulating the lipid environment *in trans* at sites of contact between organelles, an intriguing hypothesis to be evaluated in future studies.

Our study further raises the possibility that mammalian TMEM16 family proteins could have functional roles as interorganelle regulators. Indeed, mammalian TMEM16H was recently shown to regulate Ca^2+^ signaling at the ER and plasma membrane MCS^63^. Considering that mammalian TMEM16K and other family members are important modulators of cellular physiology and human pathology, determining whether and how they are involved in interorganelle communication would significantly improve our understanding of this protein superfamily as well as cellular communicome, the cellular communication network.

## Supporting information

Supplemental Table 1. TMEM16K proteomics

Supplemental Table 2. Generated constructs

Supplemental Video 1. Hindlimb clasping

Supplemental Video 2. Live imaging of TMEM16K and ER marker

Supplemental Video 3. Live imaging of TMEM16K, Rab7 and EGF

---

**Supplemental Figure 1.**

**a.** Phylogenetic tree of highly conserved family of proteins represented with yeast Ist2p, human TMEM16 family with 10 homologs (TMEM16A-H), Drosophila family with 5 homologs (Axs, Subdued, CG6938, CG10353, CG15270), fungi *Aspergillus fumigatus* homolog afTMEM16, fungi *Nectria haematococca* homolog nhTMEM16 and amoebozoa *Dictyostelium discoideum* DdTMEM16. Representation was constructed in Jalview based on Neigbourhood Joining algorithm generated from the MUSCLE alignment of the sequences. **b.** RT-PCR from liver and brain tissues obtained from the wild type and TMEM16K ubiquitous knockout littermates. RNA was extracted, reverse transcribed and equal amounts of each tissue sample were loaded on agarose gel and revealed with GelGreen. Two different sets of primers amplifying TMEM16K were used, and Gadph and β-actin were amplified as controls. DNA Ladder used was GeneRuler 1 kb Plus DNA Ladder from ThermoScientific. **c.** RT-PCR from C57BL/6J mouse tissue shows TMEM16K is broadly expressed. RNA was extracted, reverse transcribed and equal amounts of each tissue sample were loaded on agarose gel and revealed with ethidium bromide. Housekeeping gene Gapdh was used as a control.

**Supplemental Figure 2.**

**a.** Immunocytochemistry of TMEM16K tagged C-terminally with proximity biotinylation enzyme revealed with anti-HA antibody and ER-marker PDI. Scale bar 10 μm. **b.** Immunocytochemistry of TMEM16K N-terminally tagged to proximity biotinylation enzyme revealed with anti-myc antibody and ER-marker marker PDI.. Scale bar 10 μm. **c.** Immunocytochemistry evaluating retention of biotinylation activity of the BioID tagged to TMEM16K. HEK293 cells were transfected with TMEM16K tagged with biotinylation enzyme constructs and 24 hours post-transfection were exposed to no biotin, 60 min or overnight (ON) of biotin at 50 μM final concentration. Distribution of TMEM16K-mediated proximity biotinylation was revealed with fluorescently conjugated Streptavidin. Scale bar 10 μm.

**Supplemental Figure 3.**

**a.** Cellular markers evaluated in WT or TMEM16K KO cells with immunocytochemistry and confocal microscopy. Giantin and Golgin97 are Golgi markers. 2PH-PLCΔ-GFP and anti-PIP2 antibody are markers of PtdIns(4,5)P_2_. Representative images from 2 independent experiments. Scale bar 10 μm. **b.** Evaluation of 3D trans-Golgi complex morphology in the WT (n=99) and TMEM16K KO (n=82) cells by quantification of the volume, area and index of fragmentation (expressed as the TGN38 volume divided by its area for each cell) of the trans-Golgi marker TGN38 obtained from 3D reconstructions of endogenous immunolabeling using Imaris software. Data obtained from 3 independent experiments **c.** Evaluation of 3D cis-Golgi complex morphology in the WT (40) and TMEM16K KO (n=36) cells by quantification of the volume, area and index of fragmentation (expressed as volume of GM130 divided by its area for each cell) of the cis-Golgi marker GM130 obtained from 3D reconstructions of endogenous immunolabeling using Imaris software. Data obtained from 3 independent experiments. P>0.01 n.s. two-tailed Student t-test.

**Supplemental Figure 4.**

Split-GFP reconstitution assay. Immunocytochemistry of single and double transfected COS7 cells with TMEM16K-V5-GFP_11_ and tested proteins tagged with GFP_(1-10)_. Representative images from 3 independent experiments. Scale bar 10 μm.

**Supplemental Figure 5.**

**a.** Electron micrographs showing membrane contact sites between ER and endosomes in the wild type and TMEM16K knockout primary fibroblasts. Scale bar, 200 nm. The percentage of endosomes with an ER contact site defined as proximity under 30 nm was quantified. Data are from two independent experiments. (n=number of electron micrographs, n(WT)=68, n(KO)=71). P>0.01 n.s. two-tailed Student t-test. **b.** Proximity ligation assay (PLA) *in situ* evaluating the extent of ER-endosome contact sites in wild type and TMEM16K knockout primary fibroblasts. Cells were fixed, permeabilized, labelled with mouse anti-VAPB and rabbit anti-Rab7 antibodies, or with omitted primary antibodies as indicated to confirm the specificity of the observed signal, and subjected to proximity ligation assay *in situ* (PLA). Nuclei were detected with DAPI. Scale bar, 20 μm. Number of PLA puncta and nuclei from 3 independent experiments was measured, and quantified as PLA puncta per cell (n=number of non-overlapping fields of view, n(WT)=231, n(KO)=252). P>0.01 n.s. two-tailed Student t-test.

**Supplemental Figure 6.**

Immunocytochemistry to evaluate subcellular localization of TMEM16K mutants to endoplasmic reticulum. COS7 were co-transfected with ER-tdTomato labeling ER (pseudocolored in cyan) and TMEM16K mutant constructs tagged with FLAG or V5 tag, which was revealed with anti-FLAG or anti-V5 antibody, respectively (pseudocolored in magenta). Scale bar 10 μm.

**Supplemental Video 1.**

Representative hindlimb clasping of neuron specific TMEM16K WT and KO animal at 24 months of age.

**Supplemental Video 2.**

Live imaging of COS7 cells transfected with mClover3-TMEM16K (pseudocolored in magenta) and ER-tdTomato (pseudocolored in cyan). Video presents overlay of the two channels.

**Supplemental Video 3.**

Live imaging of COS7 cell transfected with TMEM17K-V5-mNeonGreen (pseudocolored in magenta), tdTomato-Rab7 (pseudocolored in yellow) and EGF-Alexa647 incubated for 45 min (pseudocolored in cyan) on spinning disk confocal. Video presents overlay of the three channels.

## Materials and methods

### Antibodies and chemicals

We used following primary antibodies: rabbit anti-V5 (Cell Signalling tech, #13202, 1/1000), mouse anti-V5 mouse (Invitrogen, # R960-25, 1/1000), mouse anti-FLAG (Sigma, # F1804, 1/1000), rabbit anti-GM130 (Abcam, # ab52649, 1/500), rabbit anti-Rab7 (Cell Signalling tech # 9367, 1/500), mouse anti-myc (kind gift from J. Michael Bishop, clone 9E10, 1/1000), rat anti-HA (Roche, # 11867431001, 1/1000), mouse anti-GFP (Roche, # 11814460001, 1/1000), rabbit anti-calreticulin (Abcam, # ab2907, 1/1000), mouse anti-PDI (Abcam, # ab2792, 1/500), rabbit anti-giantin (kind gift from Marc Von Zastrow, 1/1500), mouse anti-Golgin97 (Molecular probes, # A21270, 1/200), rabbit anti-mCherry (Abcam, # ab167453, 1/1000), chicken anti-mCherry (Novus Biologicals, #NBP2-25158, 1/200) mouse anti-EEA1 (BD Biosciences, #610456, 1/200), mouse anti-PIP2 (Abcam, # ab11039, 1/250), rabbit anti-TGN38 (Novus Biologicals, #NBP1-03495SS, 1/100), and mouse anti-VAPB (R&D systems, #MAB7329-SP, 1/100 for PLA *in situ*). Secondary antibodies (used at 1/400) and Streptavidin conjugated with Alexa Fluor 647 (# 016-600-084, 1/2000) were purchased from Jackson laboratories or Invitrogen. Phalloidin conjugated to Alexa Fluor 633 was obtained from Molecular probes (# A-22284, 1/400). Cholera Toxin Subunit B, AlexaFluor 555 conjugate was purchased form ThermoFisher (# C22843). α-Bungarotoxin, Alexa Fluor 488 conjugate (# B13422), Lysosensor Green DNP-189 (#L-7535), Alexa Fluor 647 conjugated EGF (#E-35351) and Alexa Fluor 647 conjugated transferrin (#T-23366) were purchased from Life Technologies. FCCP (#C2920-10MG) was purchased from Sigma.

### Plasmids

We obtained mouse TMEM16K cDNA from Genscript (Clone ID: OMu10422D, pCDNA3.1-TMEM16K-FLAG). Myc-BioID and BioID-HA were a gift from Kyle Roux (Addgene plasmid # 35700 and # 36047, respectively). Flag-CIMPR was a gift from Marc Von Zastrow, UCSF. CIBN-CAAX was a gift from Pietro de Camilli, Yale. GFP-P4M-SidM was a gift from Tamas Balla (Addgene, # 51469). p40PX-EGFP was a gift from Michael Yaffe (Addgene, #19010). pKanCMV-mClover3-mRuby3 was a gift from Michael Lin (Addgene plasmid # 74252). pSBtet-RB was a gift from Eric Kowarz (Addgene plasmid # 60506). pcDNA3.1-GFP_(1-10)_ was a gift from Bo Huang (Addgene plasmid # 70219). 2PH-PLCdelta-GFP was a gift from Sergio Grinstein (Addgene, # 35142). pEGFP-N1-VAPA was a gift from Axel Brunger (Addgene plasmid # 18874). tdTomato-Rab7, ER-tdTomato and mNeonGreen-mRuby2-FRET-10 were obtained from UCSF Nikon Imaging Center Library. We have obtained cDNA for VPS26A (Clone ID: BC022505) OSBPL9 (Clone ID: BC025978) and OSBPL11 (Clone ID: pCS6(BC065213)) from Transomics. We have received SNX1 cDNA as a gift from Ewan Reid, University of Cambridge. SNX2 and VPS35 cDNA were gift from Marcel Verges, Universitat de Girona. ORP8 was gift from Francesca Giordano, Institut de Biologie Integrative, Gif-sur-Yvette. mCherry-tagged GPI RUSH construct was gift from Franck Perez, Institut Curie. mCherry-tagged Transferrin Receptor (TfR) RUSH construct and GFP-tagged CD-MPR RUSH construct were gift from Juan Bonifacino, NIH.

Detailed description of all the constructs designed for this study is given in **Supplemental Table 2.** Site-directed mutagenesis was performed using PfuTurbo polymerase followed by DpnI digestion protocol. All other plasmids were constructed using classical subcloning or Gibson assembly using Hot Start High Fidelity Q5 polymerase from NEB. All constructs generated and used in this study were verified with sequencing of the coding region of the plasmids.

### TMEM16K mice models

We obtained TMEM16K conditional knockout mice (Ano10tm1a(EUCOMM)Wtsi)) generated by the International Mouse Knockout Consortium and ordered from EMMA (EMMA ID: 08927). Ubiquitous TMEM16K knockout mice were generated by crossing with the actin-driven Cre line, while crossing with nestin-Cre line generated neuron specific TMEM16K knockouts. All procedures performed have been approved by the UCSF IACUC. Wild type allele is identified with Ano10_257998_F (CACTCCCTCATCCCATTCTTG), and Ano10_257998_R (AGACGGCCACCTTACCACAG) primers (band size 433 bp). Mutant allele is detected with PCR with Ano10_257998_F and CAS_R1_Term (TCGTGGTATCGTTATGCGCC) primers (band size 156 bp). Animal cohorts were generated from heterozygote breedings, where WT and TMEM16K knockout littermates were kept gender segregated but genetically mixed with up to 5 animals per cage and aged up to 24 months.

### Neuromuscular junction staining

Animals were anesthetizes with isofluorane, and perfused first with PBS and then with 4 % PFA. In all the animals, we have dissected, analyzed and compared the same muscle localized in the hindlimbs. After isolation, muscles were immersed in blocking buffer containing 0.5% Triton-X 100 and 10% donkey serum in PBS at 4°C overnight. The muscles were then washed three times with 1 x PBS at room temperature for 15 minutes each time followed by incubation with fluorescently labeled α-Bungarotoxin overnight at 4°C on a rotating platform. The muscles are again washed 3 times with 1 x PBS, and mounted Fluoromount-G medium (SouthernBiotech) with weight applied on the cover glass to obtain flat preparation suitable for imaging. NMJ were imaged on confocal microscope as z-stacks and represented as maximum intensity projections to ensure entire structure is captured, and then analyzed in Fiji Software (ImageJ, NIH).

### Hindlimb clasping and ledge assays

Behavior of WT and TMEM16K KO littermates was analyzed using hindlimb clasping and ledge assays. Hindlimb clasping is a marker of disease progression in a number of mouse models of neurodegeneration^1^, while ledge assay is a direct measure of coordination, which is impaired in cerebellar ataxias and many other neurodegenerative disorders. The evaluation of mice was done as previously described^2^. Briefly, each measure is recorded on a scale of 0-3 depending on the severity of phenotype. For hindlimb clasping, mouse was lifted clear of all surrounding objects by grasping the tail near its base. The hindlimb position was observed for 1 min. If the hindlimbs were consistently splayed outward, away from the abdomen, it was assigned a score of 0. If one hindlimb retracted toward the abdomen for more than 50% of the time suspended, it received a score of 1. If both hindlimbs were partially retracted toward the abdomen for more than 50% of the time suspended, it received a score of 2. If its hindlimbs were entirely retracted and touching the abdomen for more than 50% of the time suspended, it received a score of 3. For the ledge assay, mouse was observed as it walked along the cage ledge and lowered itself into its cage. A wild type mouse walks along the ledge without losing its balance, and lowers itself back into the cage using its paws. This was assigned a score of 0. If the mouse lost its footing while walking along the ledge, it received a score of 1. If it did not effectively use its hind legs, or landed on its head rather than its paws when descending into the cage, it received a score of 2. If it fell off the ledge, while walking or attempting to lower itself, or shaked, it received a score of 3. All behavior analysis was done blinded of the mice genotype, with each mice tracked during the analysis with a random numeric code.

### RNA extraction and RT-PCR

Animals were euthanized with CO_2_ and tissue was immediately dissected on ice. Total RNA was extracted with TRIzol (ThermoFisher) and first-strand cDNA was synthesized with SuperScript^™^ III First-Strand Synthesis System kit (ThermoFisher) or High Capacity cDNA Reverse Transcription Kit (Applied Biosystems). We quantified the cDNA with NanoDrop and performed PCRs with Q5 Hot Start High Fidelity polymerase (NEB) using equal amounts of obtained cDNA from each tissue sample to detect presence of TMEM16K (primer set 1: CATGGCCATCATTGGACTGCCC, GCACAGCCACGCTTCCACAC, size 126 bp; primer set 2: GCCATGCGGGCCTTCACCTA, CAGTCCAATGATGGCCATGGGG, size 405 bp) and housekeeping genes Gadph (primers: TGGCCCCTCTGGAAAGCTGTG, AGTTGGGATAGGGCCTCTCTTGC, size 501 bp) and β-actin (primers: ATGAGCTGCGTGTGGCCCCTG, GACGCAGGATGGCGTGAGGG, 258 bp)

### Cell culture and transfection

Primary mouse embryonic fibroblasts (MEF) were generated from 13.5-14.5 days old embryos obtained from time pregnancies set up from heterozygote breedings. Each embryo was genotyped and individually processed for primary culture. Primary MEF culture was established using standard protocol. Primary MEF were used for maximum of 5 passages. HEK293, COS-7, U-2OS and primary MEF cells were cultured in Dulbecco’s modified Eagle’s medium (DMEM) containing 10% fetal bovine serum (FBS) and 1% penicillin/streptomycin at 37°C and 5% CO2.

We used HEK293 cells for proteomics due to their easy maintenance, suitability for scaling needed for biochemistry, and human origin to simplify peptide detection. COS-7 cells were used for imaging due to their flat morphology, ease of maintenance and transfectability. U-2OS cells were used for colocalization analysis due to their human origin suitable for larger number of antibodies and suitable flat morphology.

Transfection of plasmids into HEK293, U-2OS and COS-7 was carried out with Lipofectamine 3000 (Life Technologies), Fugene6 (Promega) or Jetprime (Polyplus transfection) following manufacturer’s instructions. Primary MEF were electroporated with the Amaxa Nucleofector using Mouse/Rat Hepatocyte Nucleofector^™^ Kit (#VPL-1004) according to manufacturer’s instructions.

The amounts of the DNA transfected were per 24-well coverslip: 20 ng for Rab7 constructs (WT Rab7, T22L, Q67N), and 500 ng for all other constructs. For electroporation we used 2 μg of DNA per construct for 2·10^6^ cells.

### Immunofluorescence

Cells were fixed for 15 min with 4% PFA; quenched for 30 min in autofluorescence reducing solution (50 mM NH_4_Cl in PBS); and blocked with 1X PBS / 5% normal donkey serum / 0.3% Triton X-100 for 30 min. Primary antibodies were incubated overnight at 4 °C in 1X PBS / 1% BSA / 0.3% Triton X-100. After 3 washes, secondary antibodies were incubated for 1 hour at room temperature before mounting in Fluoromount-G medium (SouthernBiotech) for immunochemistry, or Vectashield for 3D-SIM and histology.

### Conventional microscopy

We performed majority of the fixed and live imaging on Leica SP8-X inverted confocal system equipped with HyD hybrid detectors, adaptive focus control and Okolab environmental control incubator cage. Live imaging was performed on Nikon Ti inverted fluorescence microscope with CSU-22 spinning disk confocal, EMCCD camera, and incubator, CO_2_, and humidity control at the UCSF Nikon Imaging Center. Images are represented using pseudocolors suitable for color-blind palette.

### Evaluation of Golgi complex morphology

We analyzed Golgi complex morphology as previously reported^3^. Briefly, we performed immunochemistry on wild type and TMEM16K knockout primary cells with either antibody against TGN38 as trans-Golgi complex marker, or GM130 as cis-Golgi marker. Images were acquired on a Leica SP8-X confocal microscope, with a pinhole of 0.5 AU and voxel depth of 0.19 μm. Imaris software was used to reconstruct Golgi complex and quantify volume and area for each Golgi complex. Index of fragmentation is defined as ratio of volume and area.

### Proximity Ligation Assay (PLA) for *in situ* detection of ER-endosome contacts

Mouse wild type and TMEM16K knockout fibroblasts were seeded on poly-L-lysine-coated 8-well chamber slide at density of 20 000 cells per chamber, fixed with 4% PFA for 10min, and subjected to proximity ligation assay according to manufacturer’s protocol (SigmaAldrich, Duolink® In Situ Detection Reagents FarRed, #DUO92013). Briefly, after permeabilization with 1X PBS / 1% BSA / 0.3% Triton X-100 for 30min, the cells were subjected to blocking, incubation with mouse anti-VAPB and rabbit anti-Rab7 antibodies overnight at 4°C, hybridization with PLA probes, ligation, amplification, and mounted in Duolink mounting media with DAPI (SigmaAldrich, #DUO82040-5ML). Same procedure was done without inclusion of primary antibodies, as negative control to confirm the specificity of the observed PLA signal. Non-overlapping images were randomly acquired throughout the slide of each sample on the Leica SP8-X confocal microscope. The Fiji Software (ImageJ, NIH) was used to quantify the number of PLA puncta (https://fiji.sc) indicative of a close apposition between VAPB (ER) and Rab7 (endosome). The number of PLA puncta measured per image was expressed as ratio to number of nuclei in the same image, giving a measurement of the average number of puncta per cell.

### Electron microscopy

Primary wild type and TMEM16K knockout cells were fixed with 4% paraformaldehyde (PFA) followed by post-fixation in EM fixative (2% PFA and 2.5% glutaraldehyde in 0.1 M phosphate buffer, pH = 7.4) and processed for electron microscopy at the UCSF Veterans Affairs Medical Center Pathology Core. Electron micrographs were analyzed blinded, where endosomes and endoplasmic reticulum were separately identified and labeled using Fiji Software (ImageJ, NIH). The percentage of endosomes with an ER contact site were quantified as described previously^4^, where endosome-ER membrane contacts were defined as proximity >30 nm.

### Structured illumination microscopy (SIM)

Structured illumination microscopy (SIM) super resolution imaging was performed on DeltaVision OMX SR imaging system at the UCSF Nikon Imaging Center. We transfected U2OS cells with TMEM16K-V5, replated 6h later on high precision coverslips suitable for high-performance microscopy (Paul Marienfeld GmbH & Co, # 0117520, 1.5H, 12 mm ø), immunostained 24h later with mouse anti-V5, rabbit anti-Rab7 antibodies, and corresponding secondary antibodies, and mounted in Vectashield. Images are represented using pseudocolors suitable for color-blind palette.

### BioID proximity biotinylation

To test whether the TMEM16K BioID-fusion proteins were enzymatically active and capable of biotinylating, HEK293 cells were transfected with myc-BioID-TMEM16K or TMEM16K-BioID-HA. Biotin was added (50 μM) for 1 hour or overnight, cells were fixed, and biotinylation was visualized with a streptavidin probe conjugated to a fluorescent label.

To purify protein complexes surrounding TMEM16K via in situ BioID-catalyzed biotin labeling, we transfected HEK293 cells with BioID-myc-TMEM16K or TMEM16K-HA-BioID and incubated with 50 μM biotin overnight (12-16 hours). Mock transfected cells were processed in parallel to account for endogenously biotinylated proteins. Experiment was performed as previously described (Roux et al. 2012). Briefly, cells were washed 3x with ice cold PBS and lysed with lysis buffer with protease inhibitors (Roche). Cell lysate was centrifuged for 30 min at 15000 g and supernatant was applied to streptavidin-conjugated magnetic beads (Dynabeads® MyOne^™^ Streptavidin C1, Invitrogen, # 65001). Beads were extensively washed 5x with lysis buffer followed by 5x washes with ice-cold PBS, and flash frozen in liquid nitrogen and stored at −80°C for mass spectrometry. The experiment was performed in a minimum of three biological replicates per conditions.

### Mass spectrometry

Sample or control-incubated streptavidin magnetic beads were resuspended in 5 mM DTT in 100 mM NH_4_HCO_3_ and incubated for 30 min at room temperature. After this, iodoacetamide was added to a final concentration of 7.5 mM and samples incubated for 30 additional minutes. 0.5 μg of sequencing grade trypsin (Promega) was added to each sample and incubated at 37°C overnight. Supernatants of the beads were recovered, and beads digested again using 0.5 μg trypsin in 100 mM NH_4_HCO_3_ for 2 hrs. Peptides from both consecutive digestions were recovered by solid phase extraction using C18 ZipTips (Millipore), and resuspended in 0.1% formic acid for analysis by LC-MS/MS. Peptides resulting from trypsinization were analyzed on a QExactive Plus (Thermo Scientific), connected to a NanoAcquity^™^ Ultra Performance UPLC system (Waters). A 15-cm EasySpray C18 column (Thermo Scientific) was used to resolve peptides (90-min 2-30% gradient with 0.1% formic acid in water as mobile phase A and 0.1% formic acid in acetonitrile as mobile phase B. MS was operated in data-dependent mode to automatically switch between MS and MS/MS. The top 10 precursor ions with a charge state of 2+ or higher were fragmented by HCD. Peak lists were generated using PAVA in-house software^5^. All generated peak lists were searched against the human and mouse subsets of the SwissProt database (SwissProt.2015.12.1) (plus the corresponding randomized sequences to calculate FRD on the searches, and adding sequences for BioID when necessary), using Protein Prospector^6^. The database search was performed with the following parameters: a mass tolerance of 20 ppm for precursor masses; 30 ppm for MS/MS, cysteine carbamidomethylation as a fixed modification and acetylation of the N terminus of the protein, pyroglutamate formation from N terminal glutamine, and oxidation of methionine as variable modifications. All spectra identified as matches to peptides of a given protein were reported, and the number of spectra (Peptide Spectral Matches, PSMs) used for label free quantitation of protein abundance in the samples.

### Proteomic dataset analysis

TMEM16K proteome candidates list was generated from a minimum of 3 independent runs per condition. All proteins that had more than one peptide detected in control conditions were eliminated from further analysis. Next, only those proteins that were at least 3-fold enriched compared to control condition were considered potential interactors. We have visualized obtained candidate list as protein-protein interaction (PPI) network generated in Cytoscape^7^ using String database (Confidence score cutoff=0.7). We next analyzed network parameters with NetworkAnalyzer to obtain centrality measures and mapped size of the nodes to the degree of the node parameter, where higher degree indicates a hub. Functional enrichment of clusters in the TMEM16K PPI network were further identified and quantified (Enrichment p-value cutoff = 0.005) with String Functional Enrichment app in the Cytoscape software, where network was visualized with Edge-weight Spring Embedded Layout. Functional enrichment term corresponding to the major identified clusters were color-coded on the network. Major clusters in the TMEM16K protein interaction network were identified with MCODE cluster app in Cytoscape using default settings and represented as simplified network overlayed with corresponding labels of the previously identified functional enrichments.

### CI-MPR assay

CI-MPR assay was performed as previously reported with minor modifications^8,9^. Briefly, cells were transfected with Flag tagged CI-MPR. 24 hours later cells were starved with serum-free DMEM for 6 hours. We then incubated cells in serum free DMEM with 1/1000 mouse anti-Flag antibody (Sigma F1804) for 60 min at 37°C. They were subsequently washed with PBS, fixed with 4% PFA and then stained for the internalized antibodies by immunofluorescence. The imaging was done on confocal microscope and analyzed in Fiji Software (ImageJ, NIH). The fluorescence intensity within a 10 × 10 μm^2^ region centered on the Golgi complex was then measured. The non-Golgi vesicular fluorescence intensity was obtained by measuring the fluorescence intensity in the 10 × 10 μm^2^ region between Golgi and cell periphery. Data are presented as the non-Golgi vesicular/Golgi CI-MPR fluorescence ratio for each cell.

### Cholera toxin subunit B assay

Cells were incubated with cholera toxin subunit B (CtxB) conjugated with Alexa 555 (stock 1 mg/ml) at 1/1000 dilution in cell culture medium for 3 min at 37°C. Coverslips were washed and chased for 1 hour, washed with PBS and fixed for 15 min with 4% PFA. Cells were immunostained for Golgi marker GM130. Endosomal retrograde trafficking of CTxB was measured as amount of colocalization with the GM130 using Pearsons colocalization coefficient in Fiji Software (ImageJ, NIH).

### Transferrin internalization assay

For the pulse-chase experiment examining transferrin internalization, WT and TMEM16K KO cells were washed with ice-cold PBS and incubated at 4°C for 1 hour in DMEM containing 25 μg/ml Alexa Fluor 647-conjugated transferrin and 0.1% BSA. Unbound transferrin was removed with 2x wash with cold medium and cells were allowed to internalize the transferrin at 37°C for 1 hour. The cells were subsequently washed with cold acidic buffer (0.2 M 100% acetic acid, 0.5 M, NaCl, pH 4.2) three times to strip surface-residing transferrin and then washed with phosphate-buffered saline and fixed. Transferrin internalization was measured as fluorescence intensity of internalized transferrin per cell (A.U) in Fiji Software (ImageJ, NIH).

### EGF colocalization assay

After 16 hours of serum starvation, WT and TMEM16K KO cells were stimulated with 100 ng/ml of Alexa 647 conjugated EGF for 3 min at 37°C, and wash to remove EGF from the medium. The cells were fixed at 10 min, 15 min, 40 min and 60 min after the initial exposure to Alexa 647-EGF, and immunostained with anti-Rab7 antibody. Images were acquired on the confocal microscope and colocalization between the Alexa647-EGF and Rab7 was analyzed in Fiji Software (ImageJ, NIH) using JaCOP plugin.

### RUSH trafficking assay

The RUSH secretory assay was performed as previously published^10,11^. Briefly, WT and KO primary MEF cells were electroporated with mCherry-tagged GPI RUSH construct, mCherry-tagged Transferrin Receptor (TfR) RUSH construct or GFP-tagged CD-MPR RUSH construct. All used RUSH construct used KDEL as streptavidin hook, blocking RUSH constructs in the ER in the biotin-free medium. Biotin-free media was generated by incubating the DMEM-FBS media with streptavidin coupled to magnetic beads for 60 min. Magnetic nature of the beads allowed easy removal from the media, followed by 0.22 um filtration to ensure media sterility. Upon addition of the biotin, constructs were released from the endoplasmic reticulum in synchronized manner allowing evaluation of their transport through the biosynthetic pathway. Hence, depending on the construct used, cells were fixed at 0min, 15 min, 60 min or 120 min after biotin addition to the medium, and visualized with immunochemistry with anti-mCherry or GFP antibody. GPI is transported with a faster dynamic from the ER through the Golgi to the plasma membrane. Transferrin Receptor (TfR) is transported with a slower dynamic from the ER through the Golgi to the plasma membrane, from which it gets endocytosed and recycled back to the plasma membrane. CD-MPR is transported from the ER to the trans-Golgi, from where it bypasses plasma membrane and gets directly transported to the endosomes. The RUSH assay for different constructs is quantified on cells fixed 60 min or 120 min after biotin addition. As GPI is transported to the plasma membrane, we quantified ratio of the surface to the total GPI detected in the cell, where the amount of surface GPI was evaluated by surface staining with rabbit anti-mCherry antibody and expressed as the ratio to total GPI detected with chicken anti-mCherry antibody. In case of TfR, we evaluated Pearsons colocalization coefficient with the early endosomal marker EEA1 using JaCoP plugin in Fiji Software (ImageJ, NIH). CD-MPR transport was likewise evaluated by analyzing colocalization with the trans-Golgi marker GM130, and in comparison to mCherry-TfR RUSH construct.

### Lysosensor assay

After 16 hours of serum starvation, WT and TMEM16K KO cells were loaded for 30 min at 37°C with 1 μM Lysosensor Green DNP-189 in culture medium, washed 2x with warm live imaging medium and immediately imaged on confocal microscope for up to 3 min per coverslip to ensure comparable time loaded with the dye between experiments. For experiments where protonophore FCCP was used, time-lapse (30 second interval) was acquired and FCCP was added to final concentration of 2 μM at 120 second. Images were analyzed in Fiji Software (ImageJ, NIH) where fluorescence intensity per cell was measured.

### Protein purification

Recombinant GFP-fusion proteins were purified from BL21(DE3) cells (NEB, # C2527H) using an N-terminal H_6_-tag, as previously described^12^. Briefly, protein expression was induced at 30°C by adding 1 mM IPTG. After 2.5 hours the cells were harvested, resuspended in bacterial lysis buffer (50 mM Tris-HCl pH 7.5, 250 mM NaCl, 2 mM imidazol, 0.5 mM EDTA, 10 mM β-mercaptoethanol) and lysed with a homogeniser. After ultracentrifugation (Beckman 45 Ti rotor, 32,000 rpm, 45 minutes, 4°C) the supernatant was incubated for 1 hour at 4°C with Ni-NTA-agarose (Qiagen), washed several times with lysis buffer and the protein was eluted by incubation for 3 hours with lysis buffer containing 500 mM imidazol. Eluted protein was dialysed using Slide-A-Lyzer^™^ Dialysis Cassettes, 3.5K MWCO (ThermoFisher) against a buffer with 25 mM HEPES-KOH pH 7.4, 250 mM potassium acetate. Protein concentration was determined with Bradford assay, and proteins stored at −80°C.

### Protein Lipid Overlay Assay

Overlay assays was performed following manufacturers instructions. Nitrocellulose-immobilized phospholipids (PIP strips; Echelon Biosciences) were blocked by incubation for 1 h with 1% fatty-acid free BSA in TBST (137 mm NaCl, 2.37 mm KCl, 19 mM Tris base, 1% Tween 20). All incubations were carried out at room temperature. We incubated 10 ml of PBST supplemented with 1% fatty-acid free BSA and 10 μg of either of the purified N-terminal cytosolic domains, all of which were GFP-tagged, for 1 hour with the PIP strips. The PIP strips were washed four times with TBST and incubated for 1 hour with a 1:1000 dilution of anti-GFP antibodies in TBST supplemented with 1% fatty-acid free BSA. The membrane was again washed four times and incubated with a 1:10000 dilution of horseradish peroxidase-conjugated anti-mouse antibodies. Bound antibodies were detected by chemiluminescence with the SuperSignal West Pico Chemiluminescent Substrate (ThermoScientific, #34080) and the C-Digit Blot Scanner Imaging System (LiCor).

### Split-GFP assay

The split GFP system is based on GFP fragments containing β-strands 1-10 (GFP_1-10_) and β-strand 11 of GFP (GFP_11_) reconstituting complete β-barrel structure of GFP able to emit fluorescence when in sufficient proximity^13^. COS7 cells were transfected either with single constructs to verify their expression, localization and absence of signal in GFP channel, or double transfected to evaluate for the presence of the signal in the GFP channel. The imaging was done on Leica SP8 confocal microscope with same imaging parameters between conditions. Since TMEM16K forms a dimer^14,15^, we validated the split-GFP approach by expressing TMEM16K-GFP_1-10_ and GFP_11_-TMEM16K to reconstitute the split-GFP. We used proteins that are not considered TMEM16K interaction partners based on our proteomics, but are known to participate in similar processes/compartments (OSBPL8, OSBPL9, VPS35) as negative controls. Images are represented using pseudocolors suitable for color-blind palette.

### Statistical analysis

We used Student’s two-tailed *t*-test or single factor ANOVA with post-test Bonferroni-corrected *t*-test. We used box plot to graphically visualize data with all box-plot elements defined; the box includes the first quartile and the third quartile, with the central line representing the median. Whiskers represent the minimum and maximum values of data. X inside the box represents the mean of data. No statistical method was used to determine sample size in any of the experiments.

## Acknowledgments

We thank Matthew Klassen, Vanja Stojkovic, Caitlin O’Brien, David Crottes, Beverly Pigott and Damien Jullie for critical reading of the manuscript, and members of the Jan laboratory for discussion. We acknowledge USCF Nikon Imaging Center, Larsen Delaine and Kari Herrington for imaging support. The electron microscopy was performed by Ivy Hsien at the San Francisco Veteran Affairs Medical Center (VAMC), for which we are very thankful. This work was supported by National Institutes of Health grant (R35NS097227 and R21AG061468) to YNJ and National Institutes of Mental Health grant (R37MH0653354) to LYJ. MP was supported by a Fyssen Postdoctoral Fellowship and National Ataxia Young Investigator Award. Mass spectrometry analysis was provided by the Bio-Organic Biomedical Mass Spectrometry Resource at UCSF (A.L. Burlingame, Director) supported by the Biomedical Technology Research Centers program of the NIH National Institute of General Medical Sciences, NIH NIGMS 8P41GM103481 and NIH 1S10OD016229). YNJ and LYJ are investigators of the Howard Hughes Medical Institute.

