## Supplementary figures and images for "“TMEM16K is an interorganelle regulator of endosomal sorting”"

### Supplemental Figure 1.

**Supplemental figure 1**

**a**

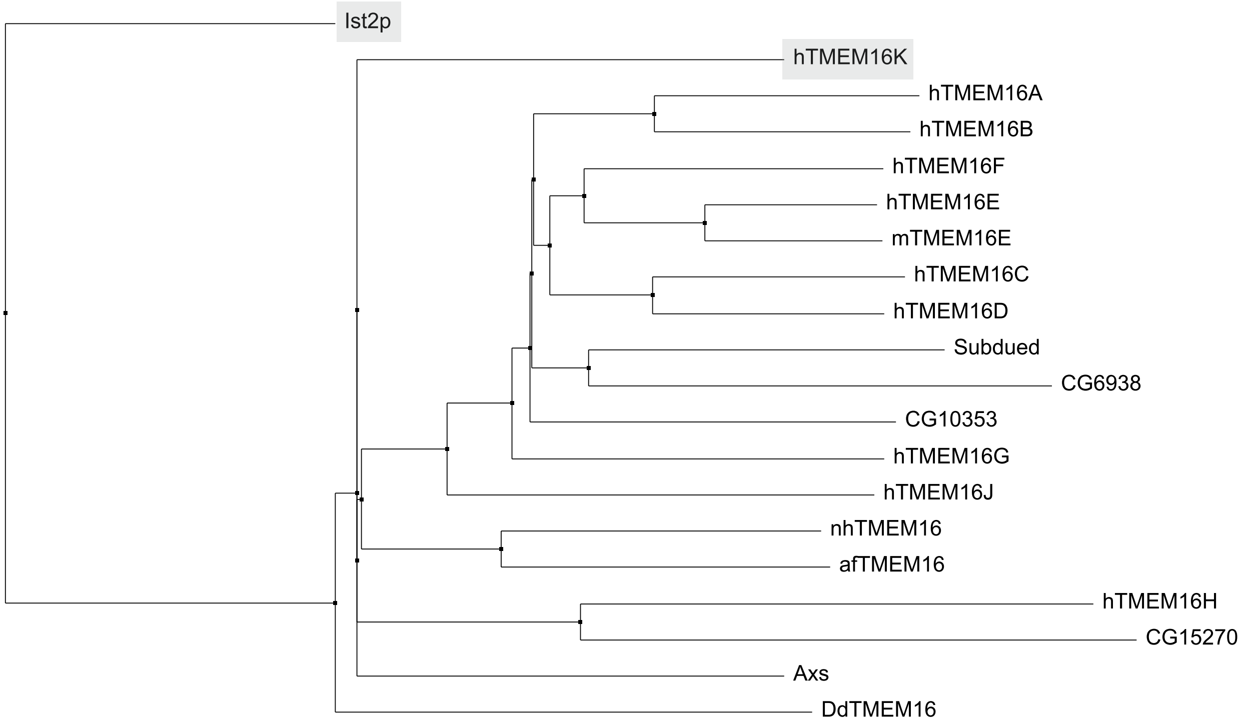


**b**

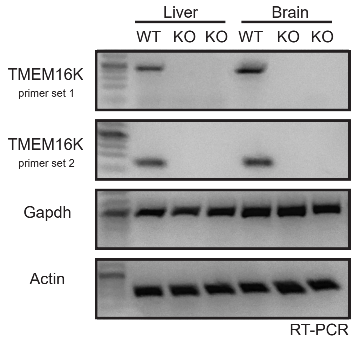


**c**

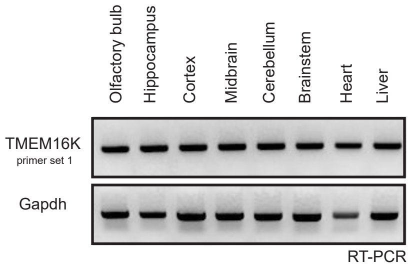

### Supplemental Figure 2.

#### Supplemental Figure 2

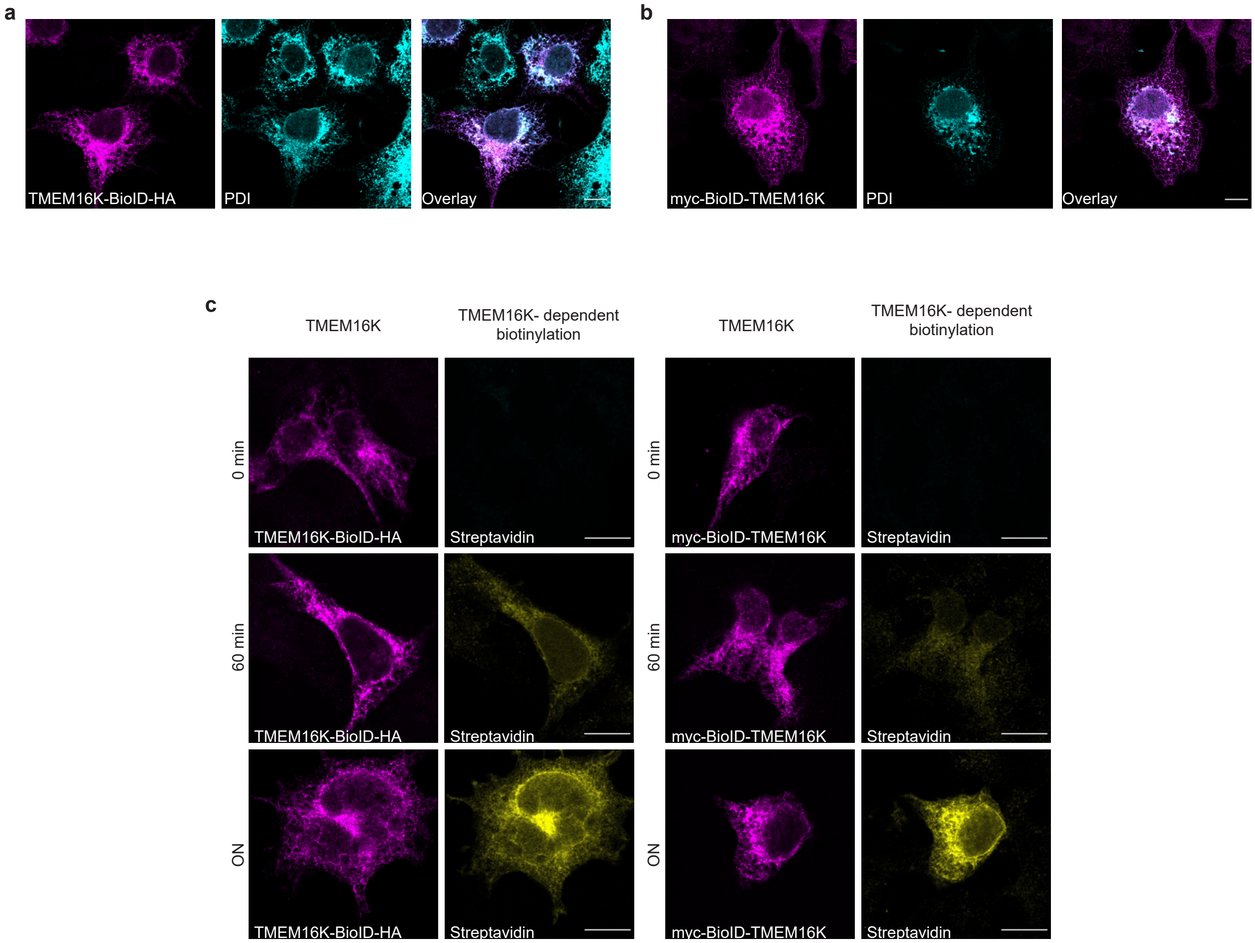

### Supplemental Figure 3.

#### Supplemental figure 3

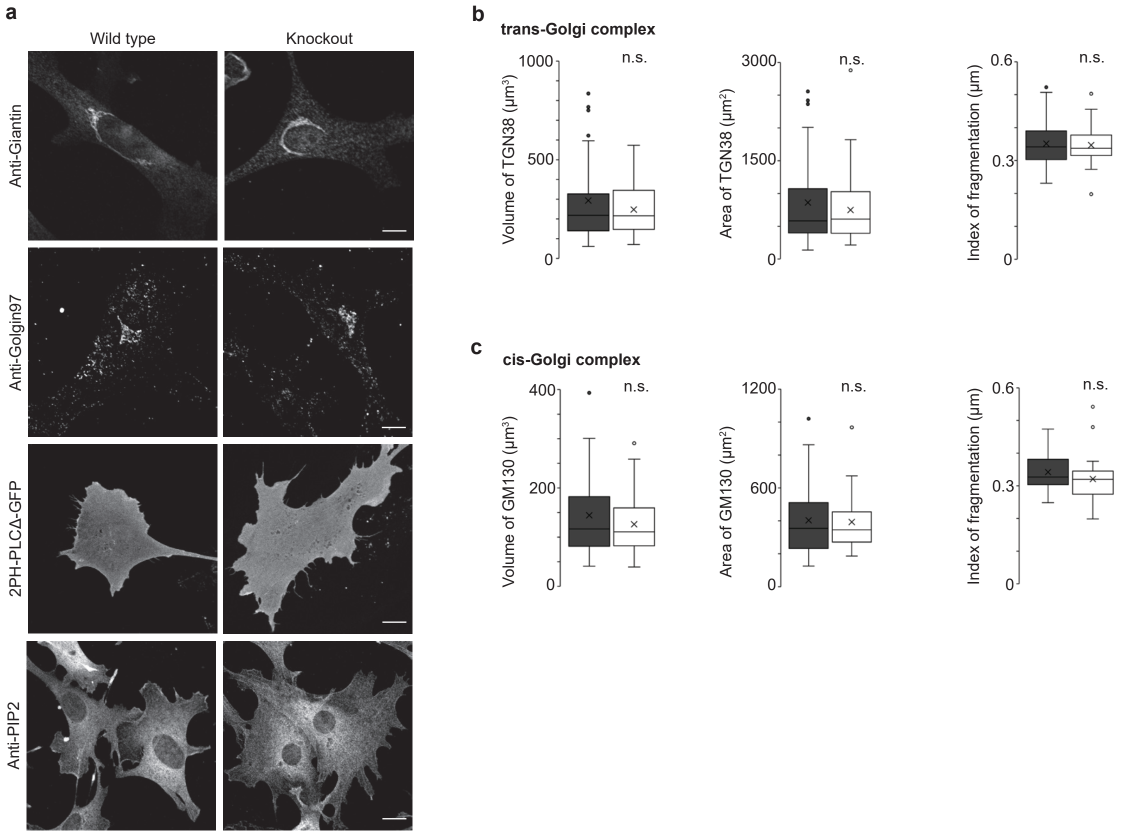

### Supplemental Figure 4.

#### Supplemental Figure 4

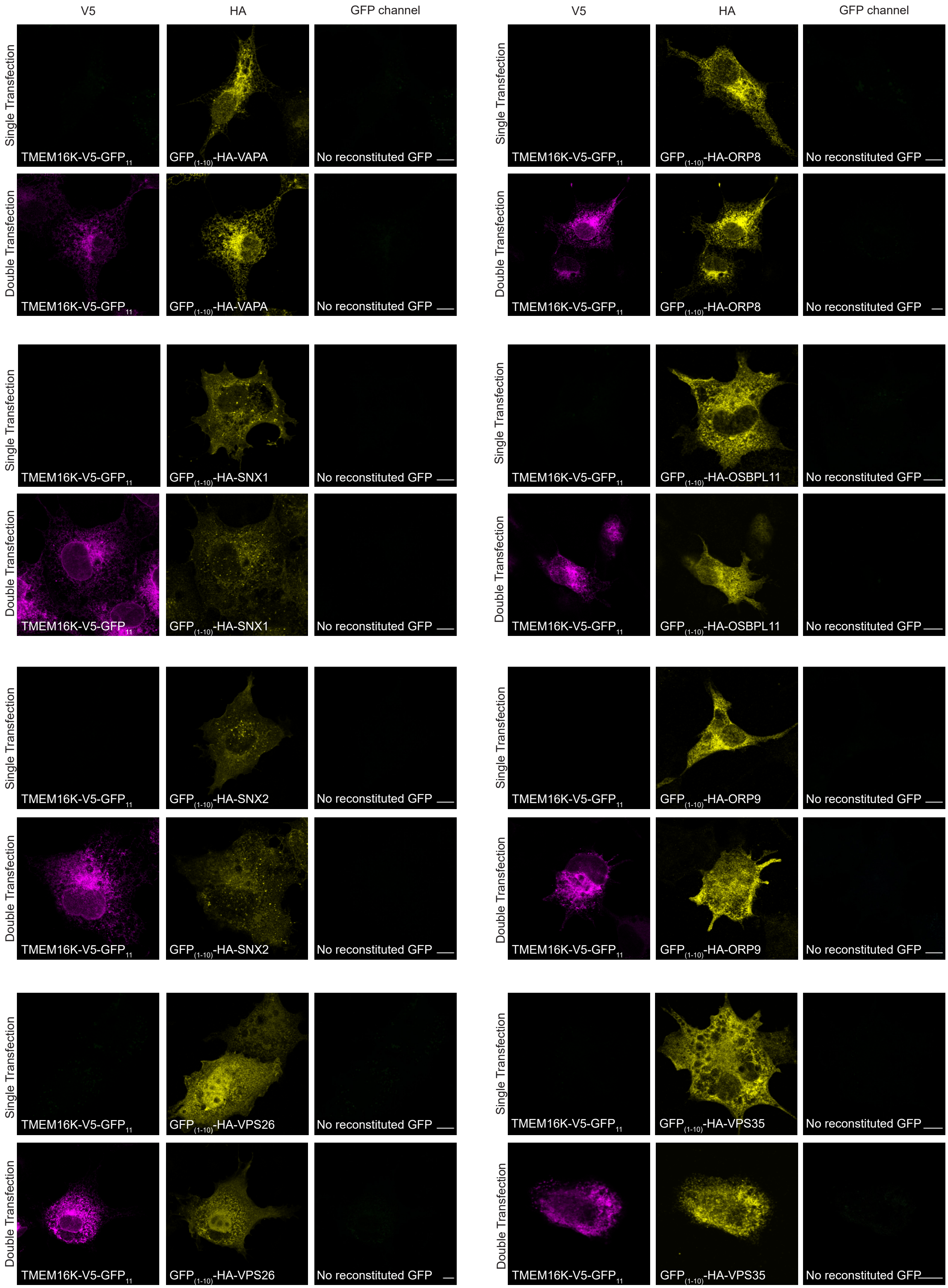

### Supplemental Figure 5.

#### Supplemental figure 5

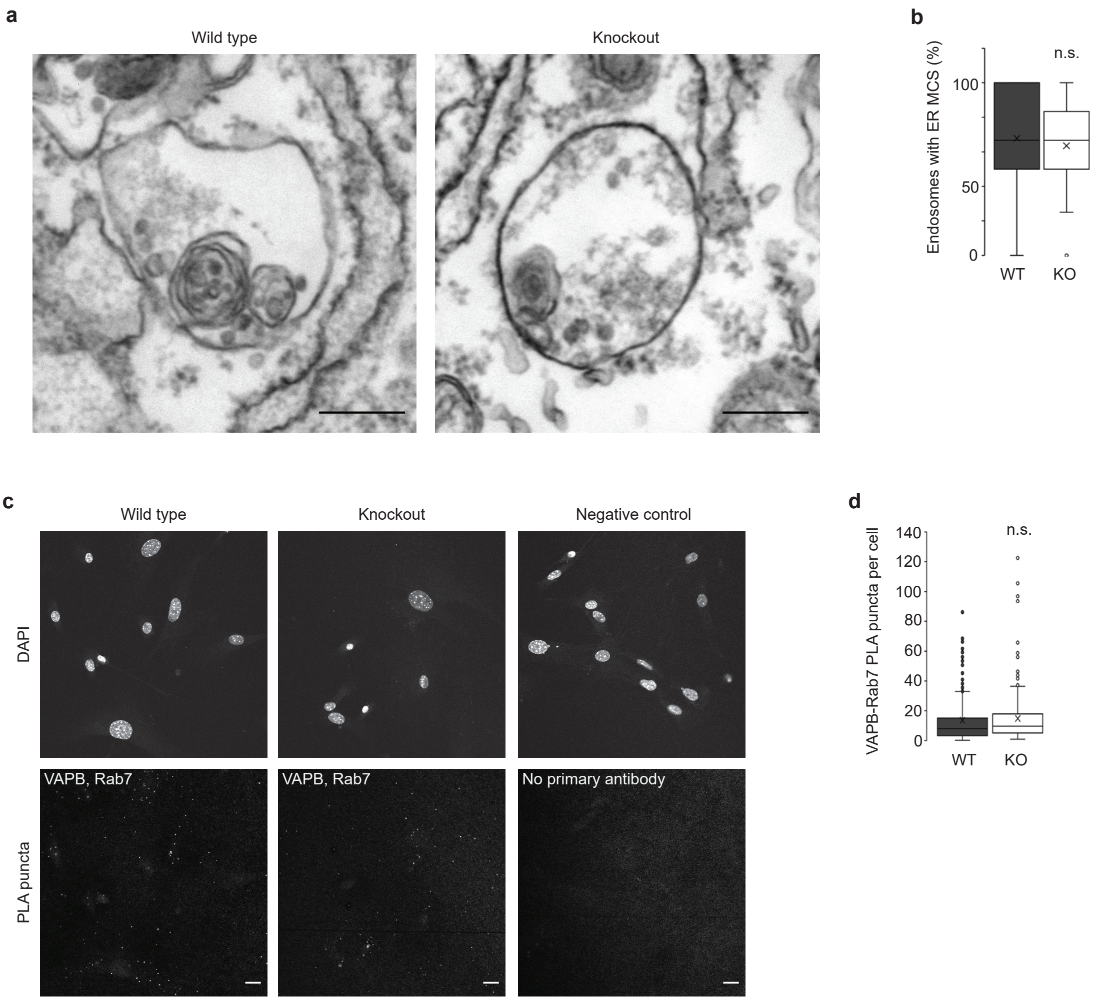

### Supplemental Figure 6.

#### Supplemental Figure 6

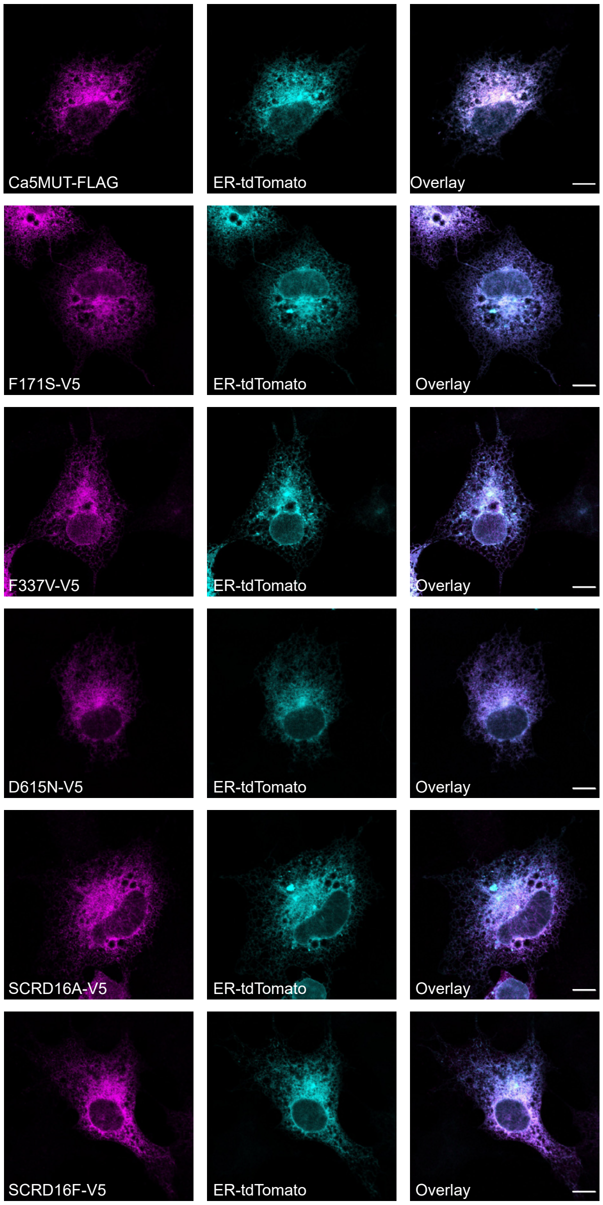
