## Supplemental Table 1. TMEM16K proteomics for "“TMEM16K is an interorganelle regulator of endosomal sorting”"

Supplementary Table 1. Candidates emerged from the proteomic mapping of protein complexes surrounding TMEM16K via *in situ* BioID-catalyzed biotin labeling

| UniProt ID | Gene Name | Peptide counts |
|---|---|---|
| Q8BH79 | TMEM16K | 219 |
| ***Endoplasmic reticulum*** | | |
| Q9BSJ8 | Esyt1 | 66 |
| Q8N766 | EMC1 | 59 |
| P51648 | ALDH3A2 | 44 |
| O95292 | VAPB | 41 |
| Q9P0L0 | VAPA | 27 |
| Q9H0U4 | Rab1b | 25 |
| Q15006 | EMC2 | 24 |
| Q92575 | Ubxn4 | 23 |
| O95573 | ACSL3 | 20 |
| Q96HY6 | DDRGK1 | 20 |
| Q9P0I2 | EMC3 | 20 |
| Q13190 | Stx5 | 15 |
| P55072 | VCP | 15 |
| Q5J8M3 | EMC4 | 14 |
| P18031 | PTP1B | 14 |
| Q9NQC3 | RTN4 | 14 |
| Q15738 | NSDHL | 13 |
| Q13586 | STIM1 | 13 |
| P27824 | CANX | 12 |
| Q96S66 | CLCC1 | 12 |
| O15260 | SURF4 | 12 |
| Q8WXH0 | SYNE2 | 12 |
| O94874 | UFL1 | 11 |
| Q8N5K1 | CISD2 | 10 |
| P61803 | DAD1 | 10 |
| P61106 | Rab14 | 10 |
| P61586 | RhoA | 10 |
| Q9NPA0 | EMC7 | 9 |
| P46977 | STT3A | 9 |
| P51572 | BCAP31 | 7 |
| Q99653 | CHP1 | 7 |
| Q9P003 | CNIH4 | 7 |
| P00387 | CYB5R3 | 7 |
| P39656 | DDOST | 7 |
| O00264 | PGRMC1 | 7 |
| Q8WVM8 | SCFD1 | 7 |
| Q5VV42 | CDKAL1 | 6 |
| Q15437 | SEC23B | 6 |
| Q9UNL2 | SSR3 | 6 |
| Q9P2W9 | Stx18 | 6 |
| Q04323 | UBXN1 | 6 |
| Q8IY17 | PNPLA6 | 5 |
| Q9NR31 | SAR1A | 5 |

| | | |
|---|---|---|
| P67812 | SEC11A | 5 |
| Q9H3N1 | TMX1 | 5 |
| Q86Y07 | VRK2 | 5 |
| O75915 | ARL6IP5 | 4 |
| Q9Y6K0 | CEPT1 | 4 |
| Q15392 | DHCR24 | 4 |
| P30519 | HMOX2 | 4 |
| Q8NF37 | LPCAT1 | 4 |
| Q99735 | MGST2 | 4 |
| P61019 | RAB2A | 4 |
| P60468 | SEC61B | 4 |
| P60059 | SEC61G | 4 |
| O00400 | SLC33A1 | 4 |
| Q9Y385 | UBE2J1 | 4 |
| Q86XL3 | ANKLE2 | 3 |
| Q07812 | BAX | 3 |
| Q6ZMG9 | CERS6 | 3 |
| Q9BXS0 | COL25A1 | 3 |
| Q9UBM7 | DHCR7 | 3 |
| Q15125 | EBP | 3 |
| Q9BW60 | ELOVL1 | 3 |
| O94905 | ERLIN2 | 3 |
| P50851 | LRBA | 3 |
| O14880 | MGST3 | 3 |
| P16333 | NCK1 | 3 |
| Q8TBX8 | PIP4K2C | 3 |
| Q92530 | PSMF1 | 3 |
| O95456 | PSMG1 | 3 |
| Q13501 | SQSTM1 | 3 |
| Q9Y3A6 | TMED5 | 3 |
| P57088 | TMEM33 | 3 |
| Q96JH7 | VCPIP1 | 3 |
| Q8WTX9 | ZDHHC1 | 3 |
| ***Endosomal transport*** | | |
| O15498 | Ykt6 | 28 |
| P51148 | Rab5c | 22 |
| Q13190 | Stx5 | 15 |
| Q96NT0 | CCDC115 | 15 |
| P46109 | CRKL | 11 |
| P61106 | Rab14 | 10 |
| P61586 | RhoA | 10 |
| O15144 | ARPC2 | 9 |
| Q9UL25 | Rab21 | 7 |
| P35232 | PHB | 7 |
| Q9BXB4 | OSBPL11 | 6 |
| P20339 | Rab5a | 6 |
| Q13596 | Snx1 | 6 |
| P51149 | Rab7a | 5 |
| Q8IWF6 | DENND6A | 4 |

| | | |
|---|---|---|
| Q9UH99 | SUN2 | 4 |
| O60784 | TOM1 | 4 |
| Q15836 | VAMP3 | 4 |
| O75436 | Vps26a | 4 |
| P50851 | LRBA | 3 |
| Q13501 | SQSTM1 | 3 |
| Q9UQN3 | CHMP2B | 3 |
| Q8IYI6 | EXOC8 | 3 |
| P30048 | PRDX3 | 3 |
| Q15286 | Rab35 | 3 |
| O14966 | Rab7L1 | 3 |
| O60749 | Snx2 | 3 |
| Q86Y82 | Stx12 | 3 |
| ***Nuclear membrane*** | | |
| P12270 | TPR | 50 |
| P46060 | RANGAP1 | 30 |
| Q5JTV8 | TOR1AIP1 | 19 |
| Q5SW79 | CEP170 | 13 |
| Q8WXH0 | SYNE2 | 12 |
| P18754 | RCC1 | 10 |
| Q9UIA9 | XPO7 | 9 |
| P49790 | NUP153 | 7 |
| Q8WUM0 | Nup133 | 6 |
| Q9UH99 | SUN2 | 4 |
| Q9Y6K0 | CEPT1 | 4 |
| P37198 | NUP62 | 4 |
| Q9BW27 | NUP85 | 4 |
| Q6ZMG9 | CERS6 | 3 |
| Q9UBM7 | DHCR7 | 3 |
| P57088 | TMEM33 | 3 |
| Q9NRG9 | AAAS | 3 |
| Q8WYP5 | AHCTF1 | 3 |
| O00629 | KPNA4 | 3 |
| O95168 | NDUFB4 | 3 |
| P35658 | NUP214 | 3 |
| P52948 | NUP98 | 3 |
| ***Proteasome*** | | |
| P43686 | PSMC4 | 9 |
| P55036 | PSMD4 | 6 |
| P17980 | PSMC3 | 5 |
| P62333 | PSMC6 | 4 |
| P60900 | PSMA6 | 3 |
| O14818 | PSMA7 | 3 |
| O00487 | PSMD14 | 3 |
| Q92530 | PSMF1 | 3 |
| ***Other*** | | |
| Q8VED5 | Krt79 | 145 |
| Q9D2U9 | Hist3h2ba | 126 |
| Q8N257 | HIST3H2BB | 126 |

| | | |
|---|---|---|
| Q6IME9 | Krt72 | 71 |
| O15173 | PGRMC2 | 50 |
| P11499 | Hsp90ab1 | 49 |
| Q9BR76 | CORO1B | 46 |
| P30041 | PRDX6 | 41 |
| Q99623 | PHB2 | 37 |
| P46821 | MAP1B | 30 |
| P50747 | HLCS | 23 |
| Q9Y266 | NUDC | 22 |
| O95721 | SNAP29 | 22 |
| P46108 | CRK | 21 |
| Q922F4 | Tubb6 | 20 |
| P07741 | APRT | 19 |
| Q00341 | HDLBP | 19 |
| Q15323 | KRT31 | 19 |
| Q14525 | KRT33B | 19 |
| P55010 | EIF5 | 18 |
| P37802 | TAGLN2 | 18 |
| Q9UPN4 | CEP131 | 16 |
| Q14247 | CTTN | 16 |
| Q9HCY8 | S100A14 | 16 |
| Q5T750 | XP32 | 16 |
| Q9H9B4 | SFXN1 | 15 |
| Q6Y7W6 | GIGYF2 | 14 |
| Q14532 | KRT32 | 14 |
| P13797 | PLS3 | 14 |
| Q9HCU5 | Sec12 | 14 |
| Q9Y490 | TLN1 | 14 |
| Q9BQ39 | DDX50 | 13 |
| Q14574 | DSC3 | 13 |
| Q9ULH7 | MRTFB | 13 |
| P08865 | RPSA | 13 |
| O75410 | TACC1 | 13 |
| Q9H2G2 | SLK | 12 |
| Q8VDJ3 | Hdlbp | 11 |
| P52732 | Kif11 | 11 |
| P08559 | PDHA1 | 11 |
| O43396 | TXNL1 | 11 |
| P30419 | NMT1 | 10 |
| O95613 | PCNT | 10 |
| P06737 | PYGL | 10 |
| Q15020 | Sart3 | 10 |
| P22735 | TGM1 | 10 |
| Q9Y277 | VDAC3 | 10 |
| O75342 | ALOX12B | 9 |
| Q9NP55 | BPIFA1 | 9 |
| Q8TDL5 | BPIFB1 | 9 |
| O75534 | CSDE1 | 9 |
| P23588 | EIF4B | 9 |

| | | |
|---|---|---|
| Q9Y2H6 | FNDC3A | 9 |
| P01876 | IGHA1 | 9 |
| Q14157 | UBAP2L | 9 |
| P08758 | ANXA5 | 8 |
| Q92747 | ARPC1A | 8 |
| Q9Y5K6 | CD2AP | 8 |
| Q9H5V9 | CXorf56 | 8 |
| Q14126 | DSG2 | 8 |
| P11171 | EPB41 | 8 |
| O76009 | KRT33A | 8 |
| P02788 | LTF | 8 |
| Q9H2M9 | RAB3GAP2 | 8 |
| Q61247 | Serpinf2 | 8 |
| P19623 | SRM | 8 |
| O95210 | STBD1 | 8 |
| P26640 | VARS | 8 |
| Q96TA2 | YME1L1 | 8 |
| Q96DA0 | ZG16B | 8 |
| Q9H3P7 | ACBD3 | 7 |
| Q96M89 | CCDC138 | 7 |
| Q8NDI1 | EHBP1 | 7 |
| P62873 | GNB1 | 7 |
| Q13765 | NACA | 7 |
| P46459 | NSF | 7 |
| Q61990 | Pcbp2 | 7 |
| Q15149 | PLEC | 7 |
| Q9H6T3 | RPAP3 | 7 |
| P16949 | STMN1 | 7 |
| Q8WUY1 | THEM6 | 7 |
| Q8N511 | TMEM199 | 7 |
| P02768 | ALB | 6 |
| Q07960 | ARHGAP1 | 6 |
| P25311 | AZGP1 | 6 |
| Q9NX63 | CHCHD3 | 6 |
| Q96C19 | EFHD2 | 6 |
| P0CG08 | GPR89B | 6 |
| O88477 | Igf2bp1 | 6 |
| Q9H910 | JPT2 | 6 |
| O76011 | KRT34 | 6 |
| P01833 | PIGR | 6 |
| P61224 | RAP1B | 6 |
| Q96FQ6 | S100A16 | 6 |
| Q15019 | SEPTIN2 | 6 |
| P28289 | TMOD1 | 6 |
| P43897 | TSFM | 6 |
| Q9Y5T5 | USP16 | 6 |
| O15143 | ARPC1B | 5 |
| Q7Z6K5 | ARPIN | 5 |
| O75964 | ATP5MG | 5 |

| | | |
|---|---|---|
| Q8WWM7 | ATXN2L | 5 |
| P35606 | COPB2 | 5 |
| Q96EB1 | ELP4 | 5 |
| Q9UBC2 | EPS15L1 | 5 |
| P11413 | G6PD | 5 |
| Q9H8Y8 | GORASP2 | 5 |
| Q9UKN8 | GTF3C4 | 5 |
| Q8N442 | GUF1 | 5 |
| P68871 | HBB | 5 |
| Q96CX2 | KCTD12 | 5 |
| Q2M1P5 | KIF7 | 5 |
| P05784 | Krt18 | 5 |
| P00846 | MT-ATP6 | 5 |
| Q9Y2A7 | NCKAP1 | 5 |
| Q9UHK0 | NUFIP1 | 5 |
| O60256 | PRPSAP2 | 5 |
| P32322 | PYCR1 | 5 |
| Q96C36 | PYCR2 | 5 |
| Q53H96 | PYCR3 | 5 |
| Q9H6Z4 | RANBP3 | 5 |
| Q5T8P6 | RBM26 | 5 |
| Q99590 | SCAF11 | 5 |
| O94901 | Sun1 | 5 |
| O43548 | TGM5 | 5 |
| Q9HAD4 | WDR41 | 5 |
| Q9NZC7 | WWOX | 5 |
| Q9NRK6 | ABCB10 | 4 |
| O75027 | ABCB7 | 4 |
| Q16864 | ATP6V1F | 4 |
| Q13867 | BLMH | 4 |
| P47755 | CAPZA2 | 4 |
| Q920C1 | Chrdl1 | 4 |
| Q86X83 | COMMD2 | 4 |
| Q9BTC0 | DIDO1 | 4 |
| Q9UGM3 | DMBT1 | 4 |
| P55039 | DRG2 | 4 |
| Q9NP97 | DYNLRB1 | 4 |
| Q16610 | ECM1 | 4 |
| Q8TE02 | ELP5 | 4 |
| Q6UN15 | FIP1L1 | 4 |
| Q14697 | GANAB | 4 |
| O75223 | GGCT | 4 |
| P09429 | HMGB1 | 4 |
| Q1KMD3 | HNRNPUL2 | 4 |
| Q14533 | KRT81 | 4 |
| P78385 | KRT83 | 4 |
| O43790 | KRT86 | 4 |
| Q9BYE3 | LCE3D | 4 |
| Q86V48 | LUZP1 | 4 |

| | | |
|---|---|---|
| Q92614 | MYO18A | 4 |
| Q13459 | MYO9B | 4 |
| P41227 | NAA10 | 4 |
| Q9BTX1 | NDC1 | 4 |
| P30414 | NKTR | 4 |
| O00151 | PDLIM1 | 4 |
| P12273 | PIP | 4 |
| Q14651 | PLS1 | 4 |
| P00491 | PNP | 4 |
| Q13136 | PPFIA1 | 4 |
| Q96QC0 | PPP1R10 | 4 |
| Q5HYI8 | RabL3 | 4 |
| O15258 | RER1 | 4 |
| O75116 | ROCK2 | 4 |
| P78345 | RPP38 | 4 |
| P06703 | S100A6 | 4 |
| Q96ES7 | SGF29 | 4 |
| P34897 | SHMT2 | 4 |
| Q8N0X7 | SPG20 | 4 |
| Q8WXA9 | SREK1 | 4 |
| P61011 | SRP54 | 4 |
| P40763 | STAT3 | 4 |
| Q6ZVM7 | TOM1L2 | 4 |
| Q9NV66 | TYW1 | 4 |
| Q14139 | UBE4A | 4 |
| Q9Y4E8 | USP15 | 4 |
| P23381 | WARS | 4 |
| Q7Z739 | YTHDF3 | 4 |
| Q96ME7 | ZNF512 | 4 |
| O00154 | ACOT7 | 3 |
| Q9H2P0 | ADNP | 3 |
| P49189 | ALDH9A1 | 3 |
| O08583 | Alyref | 3 |
| O75179 | ANKRD17 | 3 |
| O15511 | ARPC5 | 3 |
| O75787 | ATP6AP2 | 3 |
| Q9H3K6 | BOLA2 | 3 |
| P35613 | BSG | 3 |
| O96005 | CLPTM1 | 3 |
| Q9UKZ1 | CNOT11 | 3 |
| P61923 | COPZ1 | 3 |
| P13073 | COX4I1 | 3 |
| Q9BZJ0 | CRNKL1 | 3 |
| P49711 | CTCF | 3 |
| O60716 | CTNND1 | 3 |
| Q2TBE0 | CWF19L2 | 3 |
| P25685 | DNAJB1 | 3 |
| Q9BVM2 | DPCD | 3 |
| Q02487 | DSC2 | 3 |

| | | |
|---|---|---|
| Q6PDI5 | Ecpas | 3 |
| Q14232 | EIF2B1 | 3 |
| Q9NR50 | EIF2B3 | 3 |
| Q06265 | EXOSC9 | 3 |
| Q9NZB2 | FAM120A | 3 |
| O14976 | GAK | 3 |
| Q9CQQ4 | Gemin2 | 3 |
| P08754 | GNAI3 | 3 |
| P11488 | GNAT1 | 3 |
| Q8TBA6 | GOLGA5 | 3 |
| Q7Z4H7 | HAUS6 | 3 |
| O75330 | HMMR | 3 |
| O15066 | KIF3B | 3 |
| Q07866 | KLC1 | 3 |
| Q92764 | KRT35 | 3 |
| O76013 | KRT36 | 3 |
| O76015 | KRT38 | 3 |
| Q6IFX3 | Krt40 | 3 |
| Q15031 | LARS2 | 3 |
| O95202 | LETM1 | 3 |
| O95372 | LYPLA2 | 3 |
| Q9UPN3 | MACF1 | 3 |
| Q7Z7H8 | MRPL10 | 3 |
| P00395 | MT-CO1 | 3 |
| O75431 | MTX2 | 3 |
| P70670 | Naca | 3 |
| Q16795 | NDUFA9 | 3 |
| Q9UGY1 | NOL12 | 3 |
| Q76FK4 | NOL8 | 3 |
| Q9CQS2 | Nop10 | 3 |
| Q9UNZ2 | NSFL1C | 3 |
| P36639 | NUDT1 | 3 |
| Q9ULE6 | PALD1 | 3 |
| Q8NEN9 | PDZD8 | 3 |
| Q9NWS0 | PIH1D1 | 3 |
| Q10713 | PMPCA | 3 |
| O14802 | POLR3A | 3 |
| Q15181 | PPA1 | 3 |
| Q99873 | PRMT1 | 3 |
| Q96HE9 | PRR11 | 3 |
| P48634 | PRRC2A | 3 |
| Q86SE5 | RALYL | 3 |
| P43487 | RANBP1 | 3 |
| Q8IXI2 | RHOT1 | 3 |
| Q5VWQ0 | RSBN1 | 3 |
| P33764 | S100A3 | 3 |
| Q9UBE0 | SAE1 | 3 |
| Q9Y512 | SAMM50 | 3 |
| Q6PDG5 | Smarcc2 | 3 |

| | | |
|---|---|---|
| Q8TAQ2 | SMARCC2 | 3 |
| O75940 | SMNDC1 | 3 |
| P02814 | SMR3B | 3 |
| P14576 | Srp54 | 3 |
| P05455 | SSB | 3 |
| Q9UJZ1 | STOML2 | 3 |
| P53597 | SUCLG1 | 3 |
| P61956 | SUMO2 | 3 |
| Q8TC07 | TBC1D15 | 3 |
| Q8WVM0 | TFB1M | 3 |
| Q9Y2S6 | TMA7 | 3 |
| Q9JKK7 | Tmod2 | 3 |
| Q5T6F2 | UBAP2 | 3 |
| Q13404 | UBE2V1 | 3 |
| P61758 | VBP1 | 3 |
| Q9Y4P8 | WIPI2 | 3 |
| Q01831 | XPC | 3 |
| Q8WU90 | ZC3H15 | 3 |
| Q14966 | ZNF638 | 3 |
