## Supplemental Table 2. Generated constructs for "“TMEM16K is an interorganelle regulator of endosomal sorting”"

**Supplementary Table 2.** List of constructs generated for this study.

Classical subcloning

| List of constructs | Backbone | RE to cut backbone | Source of insert | Primers to amplify insert | RE to cut the insert |
|---|---|---|---|---|---|
| **myc-BioID-TMEM16K** | pcDNA3.1-myc-BioID (Addgene) | NotI, KpnI | pcDNA3.1-TMEM16K-FLAG (Genscript) | F:gcggcggccgctaagagtgactttatcaacgctggatacttgt<br>R:cccggtacctaggtagcttccttcccatcttcctggt | NotI, KpnI |
| **TMEM16K-BioID-HA** | BioID-HA (Addgene) | AgeI, BamHI | pcDNA3.1-TMEM16K-FLAG (Genscript) | F:gcgaccggtatgagagtgactttatcaacgctggatacttgt<br>R:cccggatcccaaggtagcttccttcccatcttcctggt | AgeI, BamHI |
| **mcherry-CAAX** | CIBN-CAAX (Pietro de Camilli) | NheI, AgeI | mcherry cDNA | F:ccgctagcatggtgagcaagggcgaggaggata<br>R: cgaccggtccgcttcgagatctcttgtacagctc | NheI, AgeI |
| **N16F-GFP** | pEGFP-N1 (Clontech) | NheI, EcoRI | TMEM16F-GFP (Tien et al. 2009) | F:ggcgctagcatgcagatgatgactaggaaggtcctgctgaac<br>R:gcggaattctcctgataagatccaagggctgct | NheI, EcoRI |
| **N16K-GFP** | pEGFP-N1 (Clontech) | NheI, HindIII | pcDNA3.1-TMEM16K-FLAG (Genscript) | F:gcggctagcaccgccaccatgagagtgacttttat<br>R:ggcaagctttgtctccccaaatagctacgaatgc | NheI, HindIII |
| **N16A-GFP** | pEGFP-N1 (Clontech) | NheI, EcoRI | TMEM16A-GFP (Tien et al. 2009) | F:gcggctagcccggggttgtggatggggag<br>R:ggcgaattcggccaaccttctcaccaaagta | NheI, EcoRI |
| **pet15b-His-N16F-GFP** | pet15b (Novagene, Millipore) | NdeI, BlpI | N16F-GFP | F:caagcttccatatgaccgtcagatccgctagcatgc<br>R:gatccggtgctcagcgtccatgccgagagtgatcccg | NdeI, BlpI |
| **pet15b-His-N16K-GFP** | pet15b (Novagene, Millipore) | NdeI, BlpI | N16K-GFP | F:caagcttccatatgaacaccgccaccatgagagtgact<br>R:gatccggtgctcagctcgtccatgccgagagtgatc | NdeI, BlpI |
| **pet15b-His-N16A-GFP** | pet15b (Novagene, Millipore) | NdeI, BlpI | N16A-GFP | F:caagcttccatatggttgtggatggggagcgcgag<br>R:gatccggtgctcagcgtccatgccgagagtgatcccg | NdeI, BlpI |

### Gibson assembly

| List of constructs | Backbone | Primers to amplify backbone | Source of insert | Primers to amplify insert |
|---|---|---|---|---|
| **TMEM16K-V5** | pCDNA3.1-TMEM16K-FLAG (Genescript) | Backbone is amplified in 2 segments splitting the AmpR gene (increases the probability that picked colony is a positive hit) and adding V5 tag with primers<br><u>Backbone part 1</u>:<br>AmpF: gtaagttggccgcaggttatcactcatgg<br>R1:gtccaagcccagcaaagggttaggaataggcttaccggtagcttccttcccatcttcctggtac<br><u>Backbone part 2</u>:<br>F2:cctattcctaaccctttgctgggcttggactcaacctagcgctgatcagcctcgactgt<br>AmpR: ccatgagtgataacactgcggccaacttac | V5 tag was added with primers. | |
| **TMEM16K-V5-GFP11** | pCDNA3.1-TMEM16K-FLAG (Genescript) | Backbone is amplified in 2 segments splitting the AmpR gene.<br><u>Backbone part 1</u>:<br>AmpF: gtaagttggccgcaggttatcactcatgg<br>R1: caccgcctccgccaccggtagcttccttcccatcttcctggtac<br><u>Backbone part 2</u>:<br>F2:gctgctgggattacatagcgctgatcagcctcgactgt<br>AmpR: ccatgagtgataacactgcggccaacttac | Synthesized by Integrated DNA Technologies<br>ggtggcggaggcggtggaggtaagcctattcctaaccctttgctgggcttggactcaaccggtggaggaggctcaggtggcggaggatcaggcggtggtggatcacgtgaccacatggtccttcatgagtatgtaaatgctgctgggattacatag | |
| **GFP$_{(1-10)}$-HA-TMEM16K** | pcDNA3.1-GFP$_{(1-10)}$ (Addgene) | Backbone is amplified in 2 segments splitting the AmpR gene.<br><u>Backbone part 1</u>:<br>AmpF: gtaagttggccgcaggttatcactcatgg<br>R1:gggtaacctccacctccgcctcctttttcatttggatctttgctcaggactgtttgtg<br><u>Backbone part 2</u>:<br>F2: gcttaggcggccttaagtttaaaccgc<br>AmpR: ccatgagtgataacactgcggccaacttac | pcDNA3.1-TMEM16K-FLAG (Genscript) | F:gaaaaaggaggcggaggtggaggttacccttacgatgtaccggattacgcaagagtgactttatcaacgctggatacttgt<br>R:gcggtttaaacttaaggccgcctaagctcaggtagcttccttcccatcttcctg |
| **GFP$_{(1-10)}$-HA-Rab7** | | | Rab7-tdTomato (UCSF Nikon Imaging Center) | F:gaaaaaggaggcggaggtggaggttacccttacgatgtaccggattacgcaacctctaggaagaaagtgttgctgaaggtt<br>R:gcggtttaaacttaaggccgcctaagctcagcaactgcagctttctgccg |
| **GFP$_{(1-10)}$-HA-VAPA** | | | pEGFP-N1-VAPA (Addgene) | F:gaaaaaggaggcggaggtggaggttacccttacgatgtaccggattacgcagcgaaacacgagcaaatcctggtcc<br>R:<br>gcggtttaaacttaaggccgcctaagcctacgagatgaatttccctagaaagaatccaat |
| **GFP$_{(1-10)}$-HA-SNX1** | | | SNX1 cDNA (Ewan Reid) | F:gaaaaaggaggcggaggtggaggttacccttacgatgtaccggattacgcagatccggagtcggaaggggc<br>R:gcggtttaaacttaaggccgcctaagctcaggagatggcctttgcctcag |
| **GFP$_{(1-10)}$-HA-SNX2** | | | SNX2 cDNA (Marcel Verges) | F:gaaaaaggaggcggaggtggaggttacccttacgatgtaccggattacgcagcggccgagagggaacc<br>R:<br>gcggtttaaacttaaggccgcctaagcctaggcaatggctttggcttcaggtag |
| **GFP$_{(1-10)}$-HA-VPS26** | | | VPS26A cDNA (Transomics) | F:gaaaaaggaggcggaggtggaggttacccttacgatgtaccggattacgcaatgagttttcttggaggctttttggtcca<br>R:gcggtttaaacttaaggccgcctaagctcacatttcaggctgttcggcagatgc |

| | | | | |
|---|---|---|---|---|
| **GFP$_{(1-10)}$-HA-ORP8** | | | ORP8 cDNA (Francesca Giordano) | F:gaaaaaggaggcggaggtggaggttacccttacgatgtaccggattacgcagagggaggtttggcagatggagaac<br>R:gcggtttaaacttaaggccgcctaagcctacttgaacatgaagtttattatgacttgaag |
| **GFP$_{(1-10)}$-HA-OSBPL11** | | | OSBPL11 cDNA (Transomics) | F:gaaaaaggaggcggaggtggaggttacccttacgatgtaccggattacgcacagggggggtgaaccagtgtcc<br>R:gcggtttaaacttaaggccgcctaagctcactctgctggttgtgttgttggaattatttt |
| **GFP$_{(1-10)}$-HA-VPS35** | | | VPS35 cDNA (Marcel Verges) | F:gaaaaaggaggcggaggtggaggttacccttacgatgtaccggattacgcacctacaacacagcagtcccctcag<br>R:gcggtttaaacttaaggccgcctaagcttaaaggatgagaccttcataaattggccc |
| **GFP$_{(1-10)}$-HA-ORP9** | GFP$_{(1-10)}$-HA-TMEM16K | Backbone is amplified in 2 segments splitting the AmpR gene.<br><u>Backbone part 1</u>:<br>AmpF: gtaagttggccgcaggttatcactcatgg<br>R1-ORP9: gcctcctcctccgccgcctgcgtaatccggtacatcgtaagggtaacc<br><u>Backbone part 2</u>:<br>F2-ORP9:gctgccaagcattaggcttaggcggccttaagtttaaaccgc<br>AmpR: ccatgagtgataacactgcggccaacttac | ORP9 cDNA (Transomics) | F:ggcggcggaggaggaggcatggtagaatcaattaaacactgcattgtgttgc<br>R:cttaaggccgcctaagcctaatgcttggcagcaccaagacg |
| **mClover3-TMEM16K** | pSBtet-RB (Addgene) | Backbone is amplified in 2 segments splitting the AmpR gene.<br><u>Backbone part 1</u>:<br>AmpF:gtaagttggccgagtgttatcactcatgg<br>R1:gcccttctcaccatggtggcctcagaggcctttcg<br><u>Backbone part 2</u>:<br>F2:gaaggagctacctgaggctgtcaggccaagcttc<br>AmpR:ccatgagtgatacactgcggccacttac | pKanCMV-mClover3-mRuby3 (Addgene)<br>pcDNA3.1-TMEM16K-FLAG (Genscript) | Insert 1 (mClover3):<br>F1:ctctgaggccaccatggtgagcaagggcgaggagc<br>R1:gataaagtcactctcattcttcgcctttgctcaccatcgg<br>Insert 2 (TMEM16K):<br>2F:gagcaaaggcgaagaaatagagtgactttatcaacgctggatcttgtgagag<br>2R:cttggcctgacaggctcaggtagcttccttcccatctcctg |
| **TMEM16K-V5-mNeonGreen** | TMEM16K-V5-GFP11 | Backbone is amplified in 2 segments splitting the AmpR gene.<br><u>Backbone part 1</u>:<br>AmpF:gtaagttggccgagtgttatcactcatgg<br>NG-R1: GCCCTTGCTCACCATTGAtccaccaccgccTGAtcc<br><u>Backbone part 2</u>:<br>AmpR:ccatgagtgatacactgcggccacttac<br>NG-F2: GACGAGCTGTACAAGtagcgctgatcagcctcgactgt | mNeonGreen-mRuby2-FRET-10 (UCSF Nikon Imaging Center Library) | Insert (mNeonGreen)<br>NG-F1: CAggcggtggtggaTCAATGGTGAGCAAGGGCGAGGAG<br>NG-R2: ggctgatcagcgctaCTTGTACAGCTCGTCCATGCCCATC |
| **SCRD16A-V5** | TMEM16K-V5 | Backbone is amplified in 2 segments splitting the AmpR gene.<br><u>Backbone part 1</u>:<br>R1:gccgtaaacttcatccatgatctcaatcacaacggcataaacaatgctgg<br>AmpF: gtaagttggccgcaggttatcactcatgg<br><u>Backbone part 2</u>:<br>F2: gccttcctgctcaagttcctgaattgcttcgcctcactcttctac<br>AmpR: ccatgagtgataacactgcggccaacttac | Synthesized by Integrated DNA Technologies<br>gtgattgagatcatggatgaagtttacggctgcattgccaggtggctcaccaagattgaggtcccaaagacagagaagagctttgaggagaggctaaccttcaaggccttcctgctcaagttcctgaattgcttc | |
| **SCRD-16F-V5** | TMEM16-V5 | Backbone is amplified in 2 segments splitting the AmpR gene. | Synthesized by Integrated DNA Technologies | |

| | | | | |
|---|---|---|---|---|
| | | Backbone part 1:<br>R1:ctcgtagatcgtgttcatgatctcaatcacaacggcataaacaatgctgg<br>AmpF: gtaagttggccgcaggttatcactcatg<br>Backbone part 2:<br>F2: gatgttcttgttccag ttcctgaattgcttcgcctcactcttctac<br>AmpR: ccatgagtgataacactgcggccaacttac | gtgattgagatcatgaacacgatctacgagaaggtggccatcatgatcaccaacttcgagctccca<br>aggacccagacggattatgagaacagcctgaccatgaagatgttcttgttccagttcctgaattgcttc | |
| **SCRD16A-V5-GFP11** | TMEM16K-V5-GFP11 | Backbone is amplified in 2 segments splitting the AmpR gene.<br>Backbone part 1:<br>R1:<br>CTCTCACAAGTATCCAGCGTTGATAAAGTCACTCTCAT<br>AmpF: gtaagttggccgcaggttatcactcatg<br>Backbone part 2:<br>F2: CCAGGAAGATGGGAAGGAAGCTACC<br>AmpR: ccatgagtgataacactgcggccaacttac | SCRD16A-V5 | All respective inserts were amplified with following primers<br>K-F:<br>ATGAGAGTGACTTTATCAACGCTGGATACTTGTG<br>AGAG<br>K-R: GGTAGCTTCCTTCCCATCTTCCTGG |
| **SCRD16F-V5-GFP11** | | | SCRD16F-V5 | |
| **F171S-V5-GFP11** | | | F171S-V5 | |
| **F337V-V5-GFP11** | | | F337-V5 | |
| **D615N-V5-GFP11** | | | D615N-V5 | |
| **deltaN16K-V5-GFP11** | TMEM16K-V5-GFP11 | Backbone is amplified in 2 segments splitting the AmpR gene, and removing the N-terminal domain of TMEM16K<br>Backbone part 1:<br>AmpF: gtaagttggccgcaggttatcactcatgg<br>dNtR:<br>GTCATGCAGCGGGAACACcatgccagcttgggtctccctatag<br>Backbone part 2:<br>dNtF:gacccaagctggcatgGTGTTCCCGCTGCATGACACTGA<br>AmpR: ccatgagtgataacactgcggccaacttac | | |

#### Site-directed mutagenesis

| List of constructs | Backbone | Site-directed mutagenesis primers | Comments |
|---|---|---|---|
| **Ca5MUT-FLAG (TMEM16K-E448Q/D497N/E500Q/E529Q/D533N-FLAG)** | pCDNA3.1.-TMEM16K-FLAG (Genscript) | 448-F: CTGAACCAAGTCGTAcAATCTCTTCTTCCTT<br>448-R: AAGGAAGAAGAGATTgTACGACTTGGTTCAG<br>497-500-F: CTGGGAACCTTTGATaATTACCTGcAGTTGTTCCTGCAGT<br>497-500-R: ACTGCAGGAACAACTgCAGGTAATtATCAAAGGTTCCCAG<br>529-533-F: GTTAAATAACTTCACTcAAGTCAACTCAaATGCCTTGAAAATGTG<br>529-533-R: CACATTTTCAAGGCATtTGAGTTGACTTgAGTGAAGTTATTTAAC | All 5 mutations were introduced in one multiplex-site-directed mutagenesis reaction. |
| **F171S-V5** | TMEM16K-V5 | F171S.F: GGCATCGTGACCCAAGTGTcCCCGCTGCAT<br>F171S.R: ATGCAGCGGGgACACTTGGGTCACGATGCC | |
| **F337-V5** | TMEM16K-V5 | F337V.F: TATGTCATGATGATCTACgTTGACATGGAGGACTGGG<br>F337V.R: CCCAGTCCTCCATGTCAAcGTAGATCATCATGACATA | |
| **D615N-V5** | TMEM16K-V5 | D615N.F: TCGCATTTGCCATCCCTaATAAACCACGGCACATC<br>D615N.R: GATGTGCCGTGGTTTATtAGGGATGGCAAATGCGA | |
| **GFP$_{(1-10)}$-HA-Rab7 T22N** | GFP(1-10)-HA-Rab7 | T22N-R: CTGGTTCATGAGTGAgtTCTTCCCGACTCCAGA<br>T22N-F: CTGGAGTCGGGAAGAacTCACTCATGAACCAGT | |
| **GFP$_{(1-10)}$-HA-Rab7 Q67L** | GFP(1-10)-HA-Rab7 | Q67L-R: GAGACTGGAACCGTTCCaGTCCTGCTGTGTCCC<br>Q67L-F: GGGACACAGCAGGACtGGAACGGTTCCAGTCT | |
